# Widespread oscillatory suppression in autism persists in the absence of psychotropic medication

**DOI:** 10.64898/2026.11.07.764864

**Authors:** Ana K. J. Amorim, Gustavo M. Alves, Gabriela C. L. Pontes, Luiz Dias, Beatriz M. F. Hondermann, Mariana C. C. Silva, Leandro Costa, Cecilia Ceriatte, Antonio Pereira, Laelia M. B. F. Brasil, Isabel C. N. Souza, Carla L. M. V. Machado, Bruss Lima

**Affiliations:** Centro de Medicina Tropical, Universidade Federal do Pará, Belém, Brazil; Instituto de Ciências Biológicas, Universidade Federal do Pará, Belém, Brazil; Instituto de Ciências da Saúde, Universidade Federal do Pará, Belém, Brazil; Instituto de Biofísica Carlos Chagas Filho, Universidade Federal do Rio de Janeiro, Brazil; Instituto de Tecnologia, Universidade Federal do Pará, Belém, Brazil; Centro de Atenção à Saúde da Mulher e da Crianças (CASMUC), Universidade Federal do Pará, Belém, Brazil

**Keywords:** Autism, ASD, electroencephalograpy (EEG), biomarkers for ASD, neuronal oscillations, alpha band, resting state, psychotropic medication for ASD

## Abstract

**Purpose:** Investigate the neural oscillatory activity associated with autism spectrum disorder (ASD) using electroencephalographic (EEG) recordings in children and adolescents. Compare the corresponding oscillatory power between ASD patients taking psychotropic medication with a drug-naïve group.

**Methods:** ASD diagnosis was done using DSM-IV and CARS (Childhood Autism Rating Scale). EEG recordings employed 22 scalp electrodes (10/20 system). ASD vs. neurotypical groups were matched for sex and age (5 females; average age: ∼ 10 years old). We analyzed the resting state (eyes open) EEG power spectrum during ∼ 2-minute recordings. The multitaper FFT method was used to estimate spectral power for the delta, theta, alpha, beta and gamma frequency bands. We pooled together data from several electrodes to compute comprehensive power spectra and assess the oscillatory behavior of macro brain regions.

**Results:** We observed widespread (∼4–20Hz) oscillatory suppression comprising the theta, alpha and beta frequency bands. The most widespread and robust ASD-associated power suppression was in the high theta and alpha bands. Delta and gamma bands showed no significant power change for any of the brain regions studied. Results from a subgroup of patients (N=9) who never used psychotropics confirmed these findings, indicating that they are not an artifact of medication.

**Conclusions:** Suppression in theta, alpha and beta oscillations may offer insights into the neurophysiological underpinnings of autism and could be used as an accessible and comprehensive tool in ASD diagnosis, regardless of the patients being or not on medication.

## 1 Introduction

Autism spectrum disorder (ASD) is a complex neurodevelopmental condition characterized by persistent challenges in social communication and interaction, as well as restricted interests and repetitive behaviors^1^. These features are reported in 80–90% of patients^2^ and are among the main diagnostic features of ASD^3^. The relative contribution of genetic, epigenetic and environmental factors to the etiology of ASD is still controversial^4^. Maternal infection, immune activation, and maternal autoantibodies directed towards the developing fetal brain can induce immune dysregulation and inflammation with impact on the development of the disorder^5,6^. The assumption is that the interplay between neuroinflammation and genetic factors influences normal brain development, with consequences on its morphology^7^, neurochemical balance^8^ and neural circuitry^9^.

An additional assumption is that changes in brain circuitry triggered by neuroinflammation shape neural dynamics and brain oscillations. Numerous studies have reported abnormalities in the brain oscillations of ASD patients. However, the results are notably very inconsistent across the literature. For simplicity, we will focus on the electroencephalographic (EEG) signal recorded in ASD patients during the resting state. Power in the theta band was found to increase^10–19^, decrease^20–35^ or remain unchanged^36–39^ relative to typically developing controls. The same is true for delta: increase^10,11,13,15,17–19,36^, decrease^12,22–24,27,28,30,31,35,39^ or no change^14,32,37,38^ in power. The reported behavior of higher frequency bands, such as beta and gamma oscillations, are also inconsistent. Some studies have found an increase^11–13,16,17,19,20,22,27,29,31,32,34,40^, a few have found a decrease^14,15,24,35,41–43^ and some studies have found no change^23,36–39^ in beta power relative to controls. Gamma oscillations have been associated with higher cognitive functions typically compromised in ASD patients^44,45^. However, contrary to expectation, some authors have found no average gamma oscillatory change in ASD subjects^12,32,36^, while others have found an increase^10,13,19,20,28–31,37,38,46,47^ and others a decrease^24,35,39,48^.

A prevailing theory of alpha rhythm generation is based upon thalamocortical loops of neuronal activity^49^. However, more recent findings suggest that alpha preferentially propagates from higher- to lower-order brain regions^50,51^. Alpha oscillations are believed to be negatively correlated with local cortical excitability, suggesting a suppressive role during task-irrelevant neuronal processing^52^. Additionally, alpha is thought to play a role in active task-relevant neuronal processing like attention, contextual and executive functions, and perception^53–55^. Despite this body of evidence in the neurotypical brain, reports focusing on alpha behavior in ASD patients are inconsistent. Some authors report no significant change^12,14,37–39^, while others find and increase^11,13,56–58^ and others a decrease in alpha power relative to normally developing brains^15,18,19,22–25,27–32,35,36,41,46,47,59–64^.

ASD diagnosis is based on somewhat subjective clinic evaluations that consider a wide range of signs and symptoms^1^. Concurrently, there has been an active search^33,65–67^ for more objective diagnostic methods based on the notion that ASD is associated with a distinctive neuronal dynamic. This is particularly challenging due to the heterogeneity of ASD phenotypes. Additionally, patients with autism are usually under psychotropic medication, with potential impact on precisely the brain rhythms we are trying to characterize^68^. Despite the challenges, the development of such biomarker, if reliable, could be used as a tool to address the neurophysiological underpinnings of autism. Here, we performed EEG recordings on a group of ASD and control patients matched for age and sex. Notably, a subgroup of our patients was not under any type of psychotropic medication, allowing us to characterize a drug-free ASD-associated brain dynamics. Additionally, we carried out relatively long (> 3 minutes) eyes-open resting state EEG recordings, which were submitted to multitaper spectral decomposition^69,70^. The multitaper method applied on long recording epochs allowed for reliable spectral estimates with reduced variance and high frequency resolution. We were thereby able to ascertain that autism-associated theta and alpha-band suppression, found in both medicated and non-medicated patients, is the most reliable biomarker for ASD.

## 2 Methods

### 2.1 Participants

Diagnosis for autism spectrum disorder (ASD) was carried out by a specialized medical team at the University Hospital of the Federal University of Pará, Brazil (UFPA-CASMUC) based on the DSM-IV and CARS (Childhood Autism Rating Scale). The CASMUC medical facility is part of the Brazilian public service responsible for providing free of charge healthcare to the general population. Most of the patients attended to are low-income citizens. CASMUC routinely carries out patient screening and diagnosis. After concurring with the pediatricians, we approached the family of these patients to inquire about the possibility of their child taking part in our study (Ethics Research Committee (CEP) authorization number: 20562413.0.0000.5172). This required the family to give a written informed consent for participation and to bring their child to CASMUC on a prearranged appointment to carry out the EEG recording session. The families received no payment for participating. However, we did contribute to transportation costs in a few cases. After finishing the recordings with ASD patients, we looked elsewhere (local communities, churches, etc.) for control subjects of similar socioeconomic status matched for age and sex.

Patients that showed excessive body movements during the recording sessions, a behavior usually observed in children with ASD, were excluded from analysis. Additionally, children with any history of prenatal, perinatal or neonatal complications, head injury with neurological symptoms, convulsions, epilepsy, or tics were also excluded. For those patients on medication, we asked them to interrupt the medication during the 24–48 hours prior to the EEG recording session. Our aim was to possibly reduce the impact of the medication on the underlying neuronal oscillations. However, only four patients effectively had their medications interrupted. Due to limited adhesion and the relatively short period of interruption, medication could still have a significant impact on the recorded EEG spectra. Notably, we had a subgroup of patients (N=9) that was never on psychotropic medication (Table 1). We analyzed this group separately to access the potential impact of psychotropic medication on the oscillatory behavior associated with ASD. This subgroup was compared with corresponding control subjects matched for age and sex.

**Table 1:** Subject identification (Subject ID), age, sex and medication use for all subjects studied (N = 48). Autism (Aut, N = 24) and control (Cont, N = 24) subjects matched for age and sex are depicted on the same line.

| Autism |  |  |  | Control |  |  |  |
|---|---|---|---|---|---|---|---|
| Subject ID | Age | Sex | Medication | Subject ID | Age | Sex | Medication |
| Aut-01 | 10 | male | None | Cont-01 | 10 | male | None |
| Aut-02 | 9 | male | Carbamazepine (400 mg/day) | Cont-02 | 9 | male | None |
| Aut-03 | 14 | male | Risperidone (2 mg/day) | Cont-03 | 14 | male | None |
| Aut-04 | 10 | male | Risperidone (1 mg/day) | Cont-04 | 10 | male | None |
| Aut-05 | 10 | male | None | Cont-05 | 12 | male | None |
| Aut-06 | 11 | male | Risperidone (1.5 mg/day) | Cont-06 | 10 | male | None |
| Aut-07 | 18 | male | None | Cont-07 | 18 | male | None |
| Aut-08 | 6 | male | Risperidone (3 mg/dia), Periciazine (10 mg/day) | Cont-08 | 6 | male | None |
| Aut-09 | 12 | male | None | Cont-09 | 12 | male | None |
| Aut-10 | 6 | male | Risperidone (2.25 mg/day) | Cont-10 | 7 | male | None |
| Aut-11 | 9 | male | Clomipramine (100 mg/day) | Cont-11 | 9 | male | None |
| Aut-12 | 7 | male | None | Cont-12 | 7 | male | None |
| Aut-13 | 9 | male | Risperidone (1 mg/day) | Cont-13 | 9 | male | None |
| Aut-14 | 6 | male | Risperidone (4 mg/day) | Cont-14 | 6 | male | None |
| Aut-15 | 8 | male | Risperidone (1 mg/day) | Cont-15 | 7 | male | None |
| Aut-16 | 9 | male | Risperidone (0.5 mg/day) | Cont-16 | 10 | male | None |
| Aut-17 | 16 | male | None | Cont-17 | 16 | male | None |
| Aut-18 | 8 | female | Risperidone (6 mg/day) | Cont-18 | 8 | female | None |
| Aut-19 | 10 | male | Risperidone (2 mg/day) | Cont-19 | 12 | male | None |
| Aut-20 | 9 | male | None | Cont-20 | 11 | male | None |
| Aut-21 | 26 | female | Quetiapine (400 mg/day), Escitalopram (10 | Cont-21 | 26 | female | None |
| Aut-22 | 6 | female | None | Cont-22 | 6 | female | None |
| Aut-23 | 8 | male | None | Cont-23 | 7 | male | None |
| Aut-24 | 13 | male | Risperidone (1 mg/day) | Cont-24 | 13 | male | None |

### 2.2 EEG recordings

EEG recordings were carried out during rest. No sedative-hypnotic medication was used to calm or reduce patient anxiety. The subject remained seated with their eyes open looking straight at a white wall. They were instructed to remain alert, to keep still and to relax their jaws. All subjects used ear pads to diminish auditory noise from the environment. All recordings lasted from 3–5 minutes and were carried out during a single session. From this full data segment, we extracted the central 2 minutes and 12 s segment (i.e., 132500 ms) for further analysis. Silver-silver chloride (Ag-AgCl) monopolar cup electrodes (contact surface diameter = 10 mm) with impedance below 10 kΩ.were placed on the scalp according to the 10-20 system (Jasper, 1958). We recorded from 20 standard locations, namely: frontal (F3, F4, F7, F8, Fp1, Fp2, Fz), temporal (T3, T4, T5, T6), central (C3, C4, Cz), parietal (P3, P4, Pz) and occipital (O1, O2, Oz) regions. The combined A1 and A2 activities was used as reference, while an electrode placed in the middle of the forehead was used as ground. The EEG signal was amplified 320 times, analogically filtered between 0.1 Hz (1st order, high-pass) and 100 Hz (2nd order, low-pass), digitally converted at 16-bit resolution (0.1 *μ*V/bit) and sampled at 200 samples/s using the BrainNet BNT-36 EEG recording equipment (EMSA-Medical Instruments, Brazil). An additional notch filter (40 dB attenuation) was applied at 60 Hz. EEG Screen and EEG Capture (BRAINTECH, EMSA XP, EMSA, Brazil) served as the user interface and the acquisition software, respectively.

### 2.3 Data processing

Spectral quantities were estimated for the EEG signal using the multitaper method^70^ implemented in Chronux 2.0^69^, an open-source, MATLAB-based (Mathworks Inc., Natick, USA), data analysis toolbox available at http://chronux.org. Essentially, the multitaper method attempts to reduce the variance of spectral estimates by pre-multiplying the data with several orthogonal tapers known as Slepian functions (based on Discrete Prolate Spheroidal Sequences). The frequency decomposition of multitapered data segments therefore provides a set of independent spectral estimates that, once averaged, gives rise to an ensemble estimate that is more reliable for noisy data. Mathematically, the multitapered power spectrum of a time series is defined for a given frequency as an average over all repetitions and tapers:

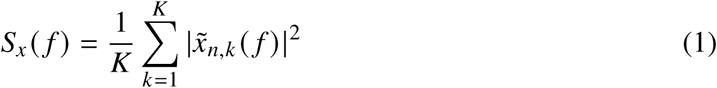

where,

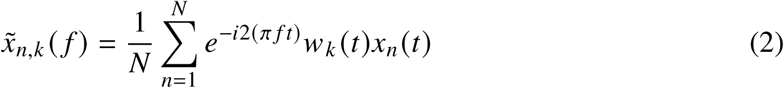

is the discrete Fourier transform of the product of the measured time series sequence {*x_n_* (*t*), *n* = 1, 2, . . . , *N*} with the k-th Slepian taper, denoted *w_k_* (*t*). Numerically, *x̃_n_*_,*k*_ ( *f*) is computed as the Fast Fourier transform (FFT) of the product. Segments of 26500 data points, (corresponding to a 2 minute and 12 s recording segment at sampling frequency of 200 points/s) were padded with zeros to the length of 131072 data points before Fourier transformation. Seventy-two Slepian tapers were used for the EEG data, resulting in a spectral concentration of ±0.54 Hz. Analyses were also computed for individual frequency bands, designated as follows: delta (1–4 Hz), theta (4–8 Hz), alpha (8–12 Hz), and beta (12–25 Hz). Regarding the gamma band, it is susceptible to ambient noise at two specific frequencies: 50 Hz and 60 Hz. The latter is directly associated with the power line, while 50 Hz noise is most probably due to equipment power supplies in the surroundings that generate noise with harmonics at 50 Hz. To minimize contamination of neuronal gamma oscillations with noise associated with these two frequencies, we discarded the full segment from 45 Hz to 75 Hz from the analysis. Gamma band was thereby defined as the segment from 25 Hz to 45 Hz, and from 75 Hz to 85 Hz. For frequency band analysis, we computed the average power within the corresponding frequency segments.

The Fourier transform was applied on a single time series (i.e., only one EEG recording was carried out per subject) and thereby resulted in a single spectrum for each EEG electrode. Here, we are interested in a comprehensive biomarker for autism. We thereby focus on the spectral behavior of entire brain regions or the entire brain. To this aim, we averaged the spectra across all electrodes of a specific brain region (occipital, parietal, central, temporal or frontal) to obtain an average estimate for each region. Alternatively, we averaged the spectra of all EEG electrodes to obtain a global estimate for the entire scalp. Finally, to obtain a population estimate, we averaged the spectra across subjects. The regional or global averages are shown separately in the results.

Overall, our data behaved as approximations to normal distributions. Relevant to us here is an easy-to-visualize comparison of the power modulation encompassing the entire 1–85 Hz frequency range between control and ASD subjects. To this aim, we used the z-statistics measure, defined as:

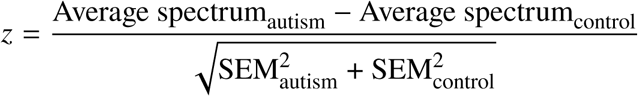

where *Average spectrum* corresponds to the *regional* or *global* average across subjects described above. The numerator represents the point-by-point subtraction of the corresponding spectra averages between autism and control, and the denominator corresponds to the square root of the sum (point-by-point) of the squared standard error of the mean (SEM) of the corresponding spectra averages. Note that despite the fact we carry out point-by-point comparisons, they involve only comparisons of the same frequency bin between spectra (i.e., they do not encompass multiple comparisons across different frequencies). Subsequently, a more rigorous power comparison between control vs. ASD conditions was done using the unpaired student t-test (see Fig. 3 and Fig. 6). In this case, we compared the average power across the entire frequency band. Significance levels were set at 95% (*p*<0.05).

## 3 Results

### 3.1 Patient characteristics

We recorded the EEG from a total of 35 patients diagnosed with ASD. Eleven of these patients showed excessive body movements during the recording sessions, a behavior usually observed in children with ASD. Muscle movements generate an EEG artifact that can be observed in the 1–85 Hz range. Critically, this frequency segment overlaps with neuronally-generated oscillatory frequencies, which are potential biomarkers for autism. Therefore, to use high quality neuronal data that could be reliably compared with sex- and aged-matched controls, these 11 patients were excluded from our analysis. This resulted in 24 ASD patients that were further analyzed (3 females, 21 males; average age 10 ± 2 standard deviation: 10.42 10 ± 2 4.53 years), for which we obtained age and gender–matched EEG recordings from control subjects (average age ± standard deviation: 10.63 ± 4.55 years).

Table 1 provides information about the age, sex and medication use by the participating subjects. Their age, represented as mean ± standard deviation (SD), were 10.62 ± 4.55 years in the control group and 10.42±4.53 years in the ASD group. Subject pairs (autism vs. control) that were matched for age and sex are depicted on the same line in Table 1. Regarding psychotropic use, 80% of the medicated patients were on risperidone, while individual patients were on either carbamazepine, periciazine, clomipramine or quetiapine + escitalopram.

Notably, we recorded from a group of ASD patients that were never submitted to psychotropic medication (i.e., the non-medicated group). This allowed us to compare the EEG results obtained from the general population (N=24) with those obtained from medication-free patients (N=9) and evaluate which modulations in the EEG spectra could be attributed to the use of psychotropic medications. The mean ages of the ASD and control subjects for the non-medicated group were 10.67 ± 4.03 years and 11.00 ± 4.09 years, respectively (*mean* ± *SD*).

### 3.2 Whole scalp and regional spectral profile of ASD and control populations

Foremost, we were interested in capturing a comprehensive whole brain oscillatory profile of our full population of ASD patients and compare it with age- and gender-matched controls. What is evident throughout our data is that autism is associated with a major suppression of brain oscillations, as captured by EEG recordings encompassing the entire scalp surface (Fig. 1). We observe a power suppression ranging approximately from 6 Hz (theta band) to 20 Hz (beta band). The alpha band, in-between the theta and beta bands, showed the most consistent suppression. We observed no frequency in which oscillatory power was enhanced in ASD as compared to control. Notably, delta and gamma power showed no significant difference between ASD and control subjects.

**Figure 1:**
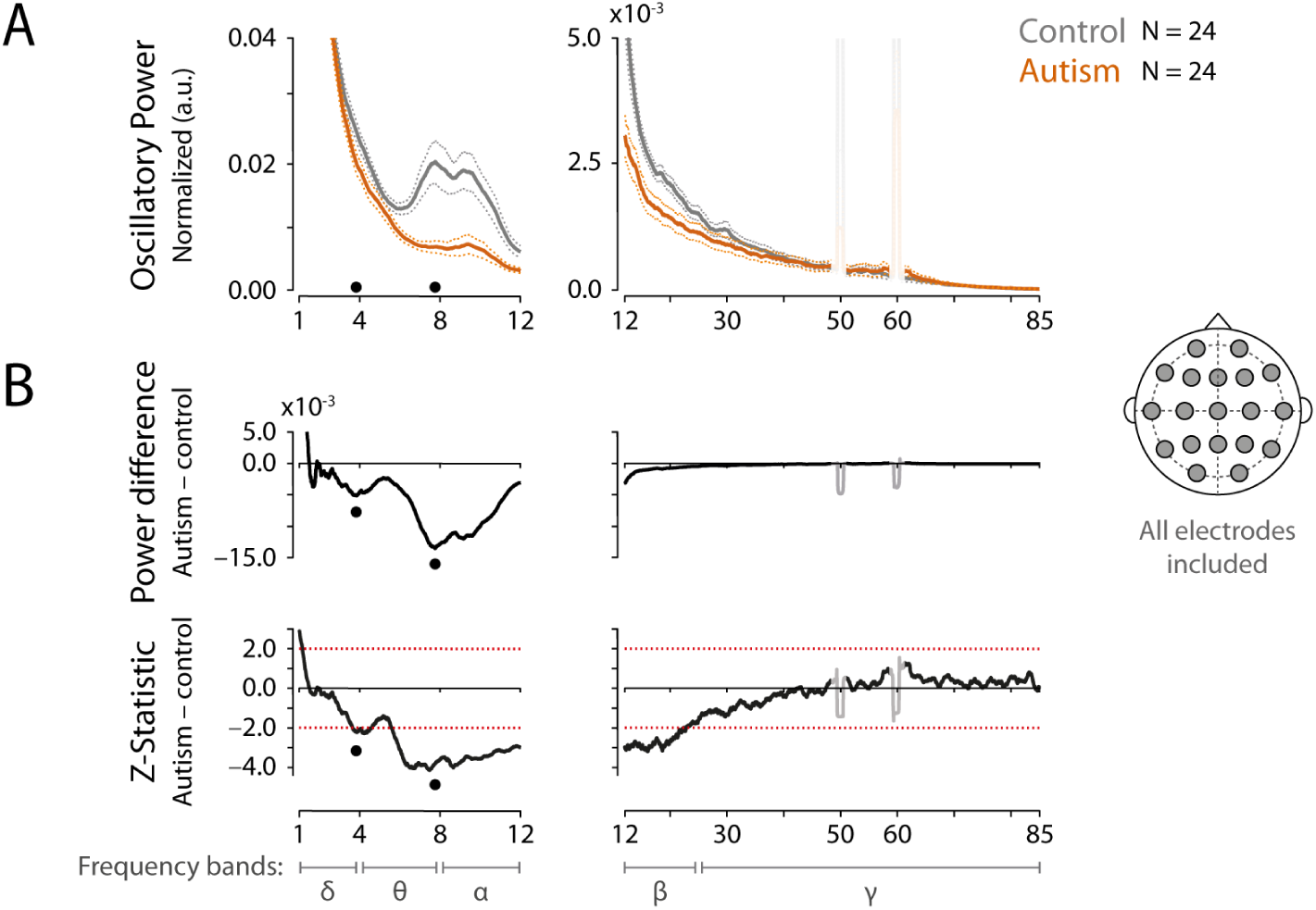
The EEG spectrum of ASD patients is associated with a pronounced suppression of alpha, theta, and beta oscillations. (A) EEG spectra computed for control (gray) or ASD (orange) subjects. For a comprehensive perspective of oscillatory behavior in autism, the average spectra (thick lines) were estimated using neural activity recorded from all subjects (N = 24) and from all EEG electrodes (see inset to the right). Thin lines correspond to the standard error of the mean (SEM). The left and right columns depict the low (1–12 Hz) and the high (12–85 Hz) frequency segments of the spectrum. (B) Top panels: Point-by-point power difference between the autism and control spectra. Black dots indicate peak differences between the spectra. To aid comparison, the black dots are mirrored on the panels above and below. Bottom panels: estimation of the Z-statistic for each point-by-point power difference (autism – control) between the spectra. Z-score values below or above −2.0 and +2.0, respectively, indicate a significant suppression or enhancement of oscillatory power (autism relative to control). *δ*, *θ*, *α*, *β*, and *γ* represent the delta (1–4 Hz), theta (4–8 Hz), alpha (8–12 Hz), beta (12–25 Hz) and gamma (25–85 Hz) frequency bands, respectively. The noise artifacts at and around 50 Hz and 60 Hz (i.e., the 45–75 Hz frequency segment) were excluded from the analysis. Note that the 1–12 Hz axis is expanded compared to the 12–85 Hz axis to provide better visualization of the power dynamics taking place at lower frequencies.

We used the multitaper method to compute all power spectra. One key advantage of this technique is that it uses multiple orthogonal tapers to concentrate the spectral energy within specific bandwidths, minimizing sidelobes due to spectral leakage. In our case, we obtained ±0.54 Hz of spectral concentration. For example, for narrowband noise peaking at 60 Hz, the multitaper technique concentrates the spectral power within a somewhat square wave ranging from approximately 59.46 Hz to 60.54 Hz (see Fig. 1). Similarly, the 50 Hz noise is concentrated within the frequency segment from approximately 49.46 Hz to 50.54 Hz. To minimize the contamination of the neural signal by these two noise sources, we discarded the full segment from 45 Hz to 75 Hz from any further analysis. Gamma band was thereby defined as the segment from 25 Hz to 45 Hz, and from 75 Hz to 85 Hz.

To further evaluate the spectral profile of ASD vs. control subjects, Fig. 2 breaks down the EEG results for the five scalp regions recorded (occipital, parietal, central, temporal and frontal regions). Spectral activity within each region was computed as the average power across all its electrodes (see inset at the bottom right of Fig. 2; activity from electrodes represented by the same color were merged). We observe the same general profile as described for the whole scalp analysis, namely a general oscillatory suppression for the joint theta, alpha and low beta segment, where the alpha band is the most consistently suppressed. Table 2 summarizes the ASD-associated power modulations (% change and associated *p* value) for each frequency band and for each brain region. Statistically significant modulations are depicted in bold. There are some particularities worth pointing out. First, even though the theta and alpha bands are both suppressed, they behave slightly different. Observe in Fig. 2A (left panels) that high theta (∼ 6–8 Hz) oscillations in control subjects are most prominent in the posterior (i.e., occipital) region of the brain and gradually decrease in power towards the frontal region. However, this same frequency segment of ∼ 6–8 Hz Hz is somewhat flat across scalp of ASD subjects. As a result, we find stronger power suppression in the occipital electrodes that gradually gets weaker towards the frontal pole (40.88% to 27.56% power decrease, respectively; see Table 2). For the alpha band, we observe the same occipital to frontal gradient in control subjects. However, the suppression seems proportional along this same posterior-to-frontal axis, averaging at around 60% of power decrease (Table 2). Second, the temporal and especially the frontal electrodes show a stronger ASD-associated suppression of the beta frequency band (Fig. 2A, top two right panels), while the occipital, parietal and central electrodes show only a modest suppression limited to the low beta band. Third, we observe no significant power differences between control and ASD subjects for frequencies below 4 Hz (i.e., delta band) or above 25 Hz (i.e., gamma band).

**Figure 2:**
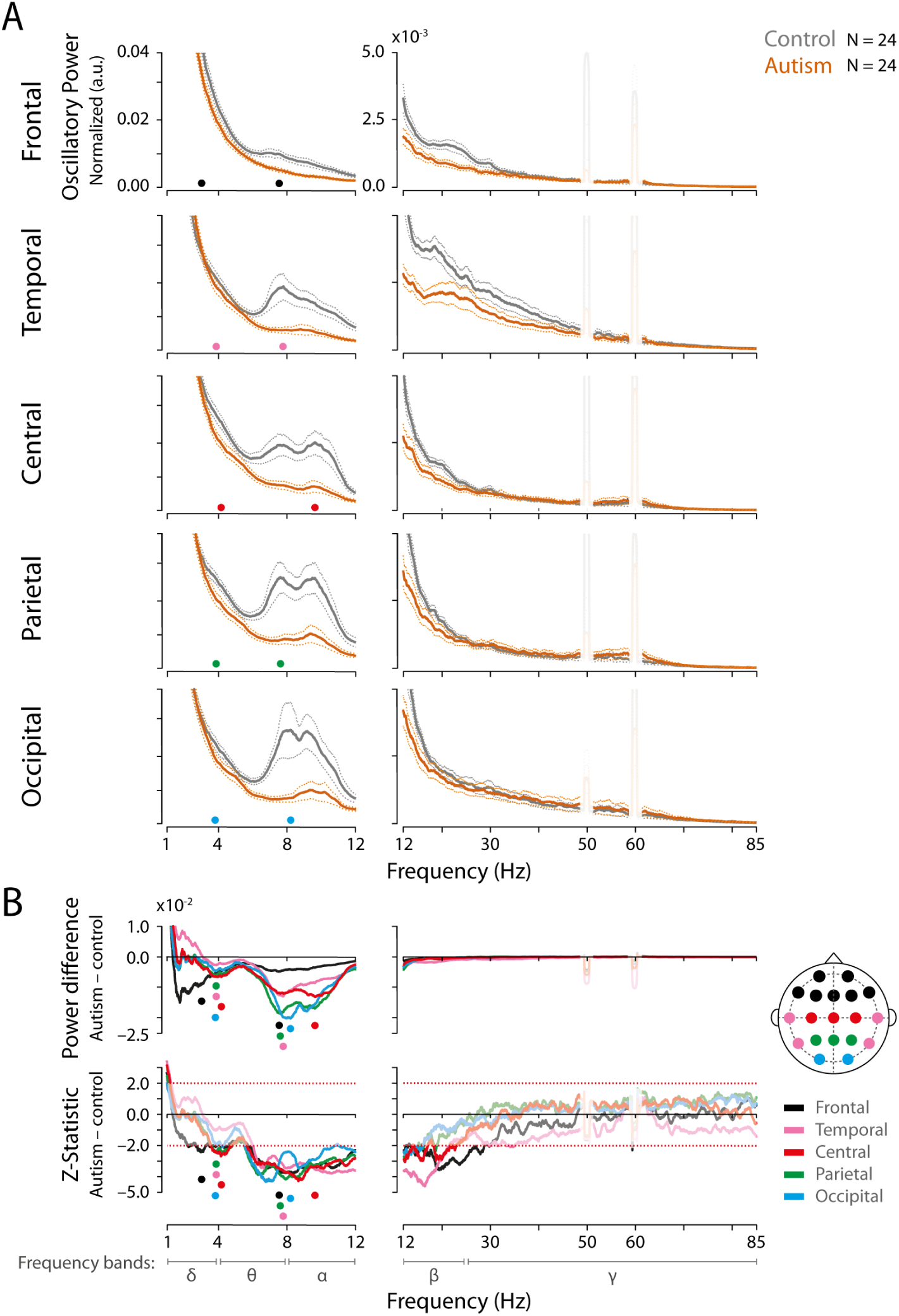
Autism-associated alpha and theta oscillatory suppression are observed in all brain regions recorded. This figure follows the same conventions as in Figure 1 but broken down for the five brain regions studied (i.e., occipital, parietal, central, temporal and frontal regions). (A) To allow for a comprehensive perspective of oscillatory behavior in autism, the average spectra (thick lines) were estimated using the pool activity from all electrodes belonging to each of the five brain regions studied and subsequently pooled across all subjects (N=24). Thin lines correspond to the standard error of the mean (SEM). Control and ASD subjects are depicted in gray and orange, respectively. (B) Top panels: Point-by-point power difference between autism and control. The five brain regions are represented by different colors (see inset to the right). Peak differences are indicated by the corresponding color dots, which are mirrored on the panels above and below to aid comparison. Bottom panels: estimation of the Z-statistic for each point-by-point power difference (autism – control) between the spectra. Z-score values below or above –2.0 and +2.0, respectively, indicate a significant suppression or enhancement of oscillatory power (autism relative to control). *δ*, *θ*, *α*, *β* and *γ* represent the delta (1 – 4 Hz), theta (4 – 8 Hz), alpha (8 – 12 Hz), beta (12 – 25 Hz) and gamma (25 – 85 Hz) frequency bands, respectively. The noise artifacts at and around 50 Hz and 60 Hz (i.e., the 45 – 75 Hz frequency segment) were excluded from the analysis.

**Table 2:** Percentage change (% change) of EEG spectral power (autism relative to control) and its corresponding statistical significance (*p* value) for 5 frequency bands (delta, theta, alpha, beta and gamma) and 5 brain regions (frontal, temporal, central, parietal and occipital). Data from all subjects (ASD: N = 24, Control = 24). The combined activity recorded from all electrodes (’All regions’) is also shown. Statistically significant changes are depicted in bold.

|  | **Frequency Band** |  |  |  |  |  |  |  |  |  |
|---|---|---|---|---|---|---|---|---|---|---|
|  | **Delta** |  | **Theta** |  | **Alpha** |  | **Beta** |  | **Gamma** |  |
| **Brain region** | **% change** | ***p*** | **% change** | ***p*** | **% change** | ***p*** | **% change** | ***p*** | **% change** | ***p*** |
| All regions | −8.59 | 0.50 | **−36.56** | **3.9 x 10<sup>-4</sup>** | **−60.69** | **3.8 x 10<sup>-5</sup>** | **−31.74** | **7.6 x 10<sup>-3</sup>** | −11.45 | 0.56 |
| Frontal | −14.92 | 0.17 | **−27.56** | **5.5 x 10<sup>-3</sup>** | **−50.52** | **8.0 x 10<sup>-4</sup>** | **−36.98** | **7.4 x 10<sup>-3</sup>** | −23.25 | 0.21 |
| Temporal | +2.58 | 0.87 | **−35.21** | **9.0 x 10<sup>-3</sup>** | **−61.24** | **1.6 x 10<sup>-4</sup>** | **−40.08** | **1.9 x 10<sup>-3</sup>** | −31.10 | 0.15 |
| Central | −6.34 | 0.60 | **−34.69** | **5.1 x 10<sup>-3</sup>** | **−62.30** | **2.9 x 10<sup>-5</sup>** | **−30.17** | **1.8 x 10<sup>-2</sup>** | +3.61 | 0.86 |
| Parietal | −10.01 | 0.47 | **−40.91** | **7.8 x 10<sup>-4</sup>** | **−60.90** | **8.8 x 10<sup>-5</sup>** | −27.89 | 0.07 | +11.91 | 0.65 |
| Occipital | −11.86 | 0.43 | **−40.88** | **1.3 x 10<sup>-4</sup>** | **−61.94** | **6.3 x 10<sup>-4</sup>** | −22.28 | 0.10 | +5.89 | 0.84 |

A population analysis based on averages across entire frequency bands (delta, theta, alpha, beta and gamma) is shown in Fig. 3. First, we show the power averages computed from the activity acquired from all EEG electrodes (Fig. 3A). Circles represent individual subjects within our entire population (N=24). When all electrodes are jointly analyzed, clear ASD-associated power suppressions are observed for the theta, alpha and beta bands (Fig. 3A). No significant differences are seen for the delta and gamma bands when we compare control vs. autism (we use pale colors to denote comparisons where *p*>0.05). In Fig. 3B, we break down the oscillatory behavior for the five brain regions (occipital, parietal, central, temporal and frontal), focusing on the three frequency bands that are significantly modulated in ASD: theta, alpha and beta. Theta and alpha are consistently suppressed in all regions, while beta oscillations are only affected in the more anterior regions of the brain (central, temporal and frontal).

**Figure 3:**
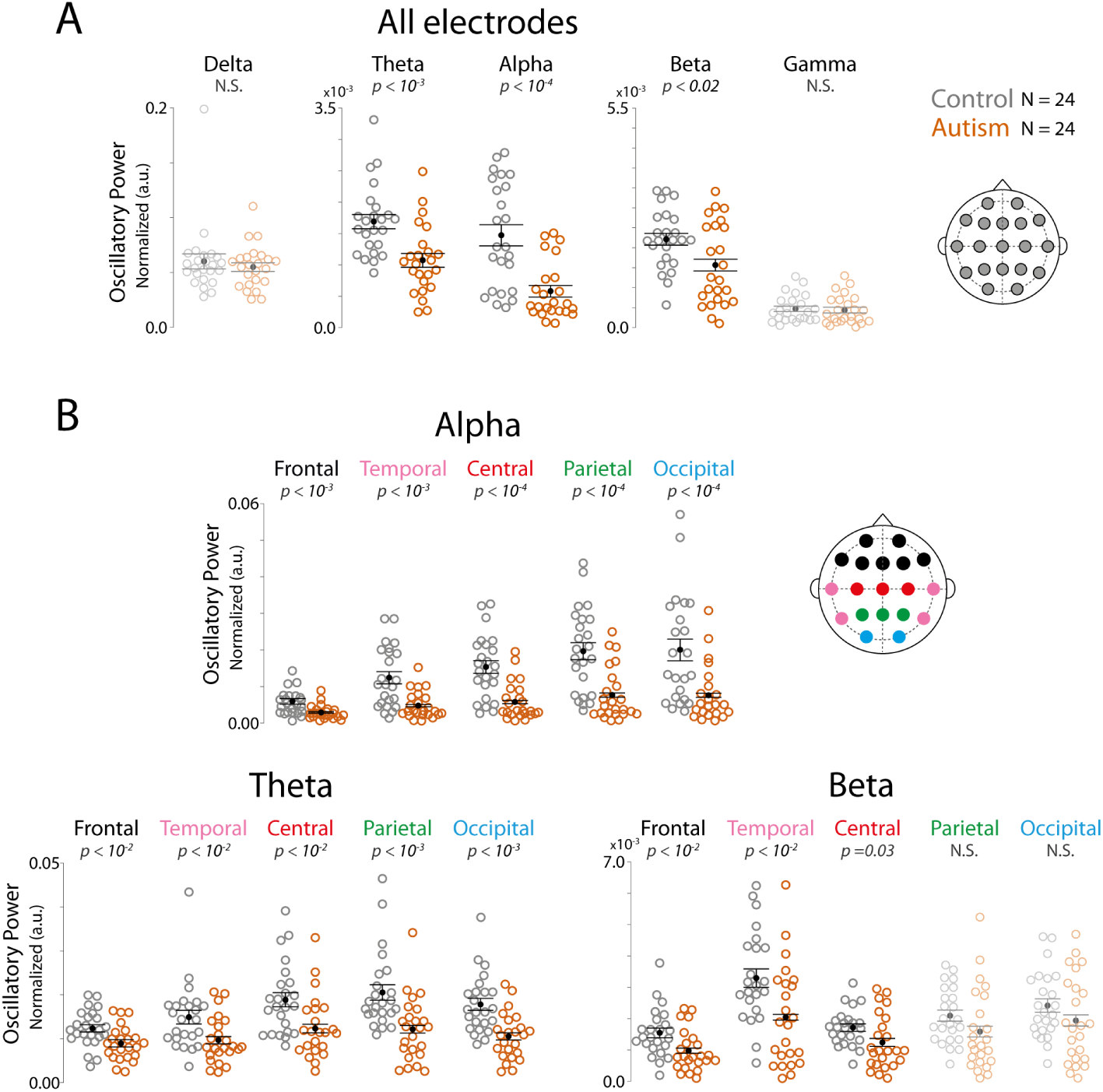
Full-band spectral power comparisons corroborate alpha, theta, and beta suppressions as biomarkers for ASD. (A) Open circles represent the spectral power averaged within a certain frequency band for each subject across all electrodes. Smaller filled black dots represent the spectral power further averaged across all subjects. The associated thin line segments correspond to the standard errors of the mean (SEM). Control and ASD subjects are shown in gray and orange, respectively (N = 24 each). Control vs. autism pairs exhibiting significant differences in their means (*p* < 0.05) are highlighted, while those pairs that do not exhibit a significant difference are shown in pale colors. (B) Same conventions as in Panel A, but the results are broken down for the five brain regions studied (i.e., occipital, parietal, central, temporal and frontal regions). Results for the delta and gamma bands are not shown because they yielded no significant differences between control vs. ASD (see Figs 1 and 2). The frequency bands were defined as follows: delta (1–4 Hz), theta (4–8 Hz), alpha (8–12 Hz), beta (12–25 Hz) and gamma (25–85 Hz). The noise artifacts at and around 50 Hz and 60 Hz (i.e., the 45–75 Hz frequency segment) were excluded from the analysis.

Figure 1, 2 and 3 capture the spectral behavior of all our ASD subjects (N = 24), whether they are under psychotropic medication (N = 15) or not (N = 9). However, a valid concern is whether the effects we describe above can be partially or totally accounted for by the medication they were taking. We were able to address this concern by carrying out the same analysis shown in Figure 1, Figure 2 and Figure 3, but restricted to the subgroup of ASD patients that had never been prescribed any psychotropic medication (Fig. 4, Fig. 5, and Fig. 6, respectively). First, we compare the average spectrum computed from all scalp recordings between non-medicated ASD subjects and their age- and gender-matched controls. As shown in Figure Fig. 4, we obtain equivalent results to those observed for the entire group (compare with Fig. 1), namely a power suppression encompassing oscillations in the theta, alpha and beta frequency bands. Fig. 5A shows that when we break down this analysis for the individual brain regions, we continue to observe power suppression in the theta-alpha-beta range, but with some features worth noting. First, in non-medicated ASD subjects, theta suppression is absent in the frontal region (see Fig. 5B, bottom panel and Fig. 6B, bottom left panel). For the central and temporal regions, we only observe significant suppression for the higher theta band (i.e., >∼ 7 Hz). Critically, we see no power suppression at the beginning of the theta band (∼ 4 Hz) for any of the brain regions (Fig. 5B), even though we observe ASD-associated suppression when the entire group of subjects is analyzed (see Fig. 1B and Fig. 2B, bottom panels, at ∼ 4 Hz). Future experiments, which include a larger population of subjects will be required to better access if psychotropic medication, as the ones used in this study (mainly risperidone) has an effect in suppressing power in the low theta band.

**Figure 4:**
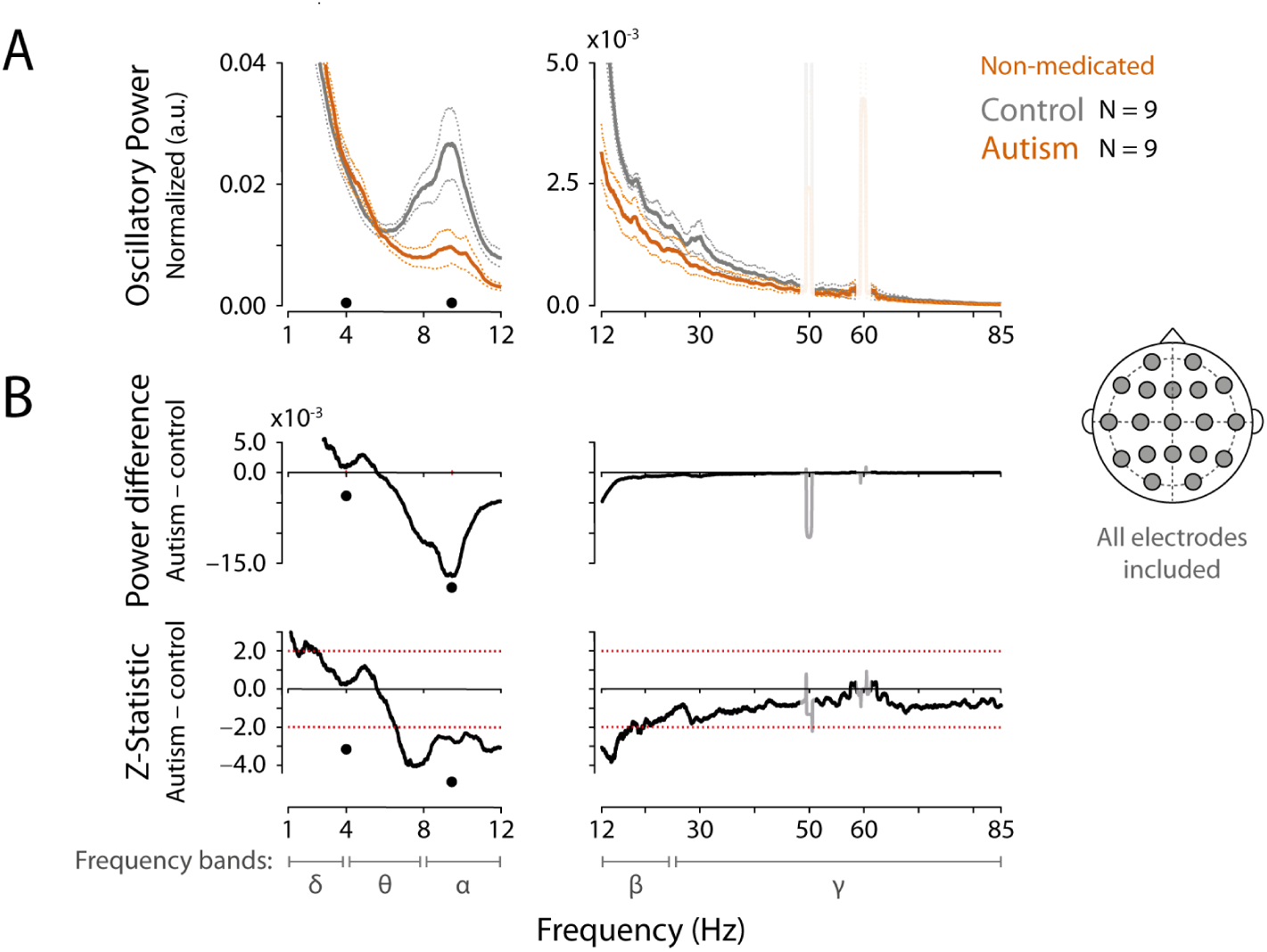
Recordings from ASD patients not taking any psychotropic medication (n=9) corroborate the results obtained with our full population (N=24). Same conventions as in Fig. 1. (A) EEG spectra computed for control (gray) or ASD (orange) subjects. The average spectra (thick lines) were estimated by pooling neural activity recorded from all non-medicated subjects (N=9) and from all EEG electrodes (see inset to the right). Thin lines correspond to the standard error of the mean (SEM). The left and right columns depict the low (1 Hz–12 Hz) and the high (12 Hz–85 Hz) frequency segments of the spectrum. (B) Top panels: Point-by-point power difference between the autism and control spectra. Black dots indicate peak differences between the spectra. Black dots are mirrored on the panels above and below (to aid comparison). Bottom panels: estimation of the Z-statistic for each point-by-point power difference (autism − control) between the spectra. Z-score values below or above −2.0 and +2.0, respectively, indicate a significant suppression or enhancement of oscillatory power (autism relative to control). *δ*, *θ*, *α*, *β* and *γ* represent the delta (1–4 Hz), theta (4–8 Hz), alpha (8–12 Hz), beta (12–25 Hz) and gamma (25–85 Hz) frequency bands, respectively. The noise artifacts at and around 50 Hz and 60 Hz (i.e., the 45–75 Hz frequency segment) were excluded from the analysis.

**Figure 5:**
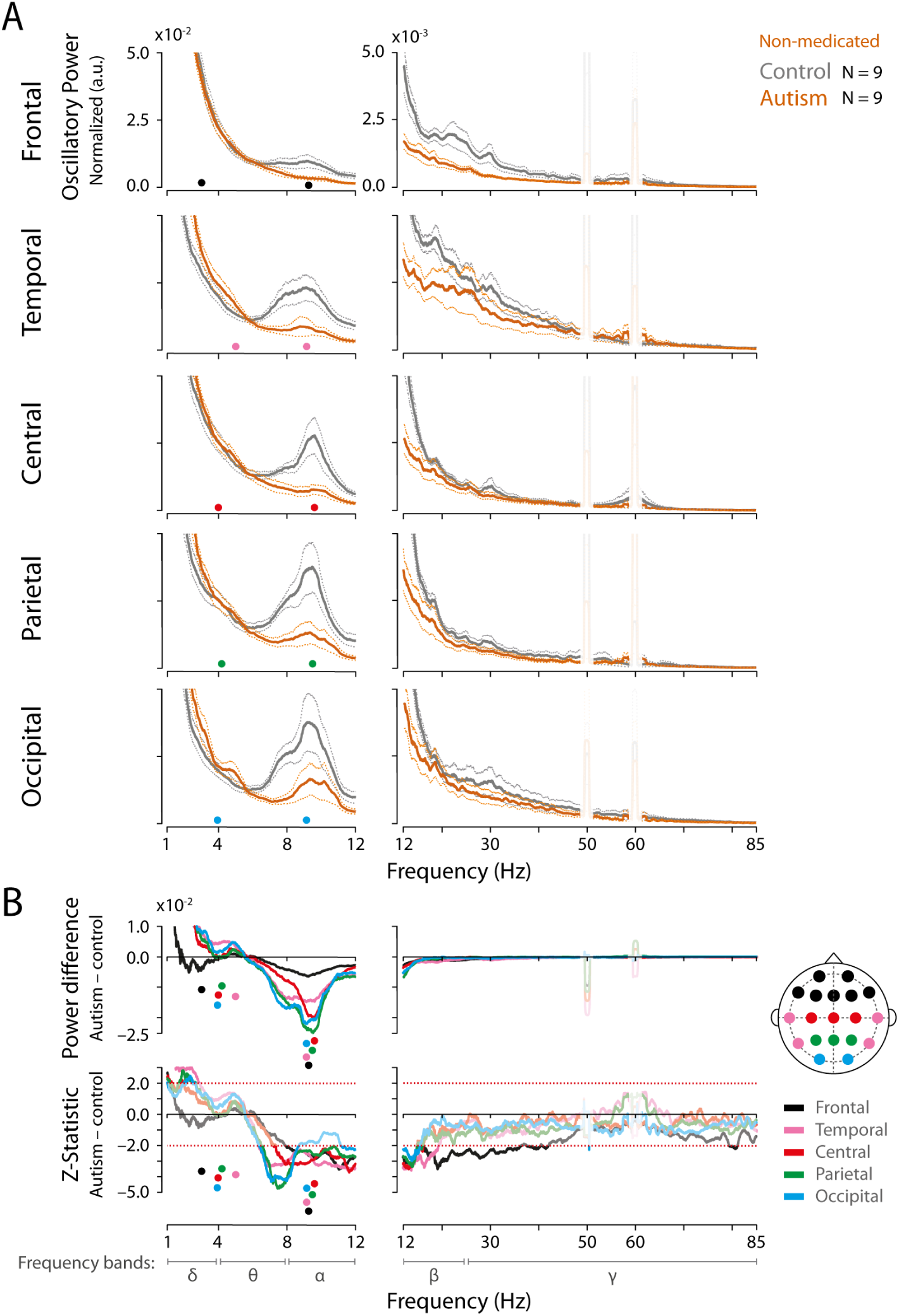
Recordings from individual brain regions of ASD patients not taking any psychotropic medication corroborate our previous results for the full population. Same conventions as in Fig. 2. The spectra were estimated using the pool activity from all electrodes belonging to each of the five brain regions studied (i.e., occipital, parietal, central, temporal and frontal regions; see inset to the right) and subsequently averaged across all subjects (N=9). (A). Control and ASD subjects are depicted in grays and orange, respectively. Thin lines correspond to the standard error of the mean (SEM). The left and right columns depict the low (1–12 Hz) and the high (12–85 Hz) frequency segments of the spectrum. (B) Top panels: Point-by-point power difference between autism and control. The five brain regions are represented by different colors (see inset to the right). Peak differences are indicated by the corresponding color dots, which are mirrored on the panels above and below to aid comparison. Bottom panels: estimation of the Z-statistic for each point-by-point power difference (autism − control) between the spectra. Z-score values below or above −2.0 and +2.0, respectively, indicate a significant suppression or enhancement of oscillatory power (autism relative to control). *δ*, *θ*, *α*, *β* and *γ* represent the delta (1–4 Hz), theta (4–8 Hz), alpha (8–12 Hz), beta (12–25 Hz) and gamma (25–85 Hz) frequency bands, respectively. The noise artifacts at and around 50 Hz and 60 Hz (i.e., the 45–75 Hz frequency segment) were excluded from the analysis.

Second, ASD-associated beta suppression continues to be prevalent at more anterior regions in the scalp. This can be better observed in Fig. 6B, which shows the population analysis for non-medicated subjects based on the average power across entire frequency bands. Here, we observe a clear suppression of frontal beta power, as corroborated by the spectral analysis shown in Fig. 5A (top right panel). Suppression for the temporal region is also visible (panel below), albeit with higher variance. Suppression at more posterior regions (central and parietal) is weaker but still significant. Therefore, we can clearly point out in our data an anterior-to-posterior ASD-associated gradient of beta suppression. The opposite is true for the theta band (Fig. 6B, where we observe a posterior-to-anterior gradient in theta suppression. Finally, the alpha band continues to be suppressed in all brain regions of non-medicated patients (Fig. 5 and Fig. 6). The magnitude of the suppression, as observed for the entire group of patients, remains estimated at around 60% (Table 3). However, the significance of the effect is weaker due to the smaller number of subjects.

**Figure 6:**
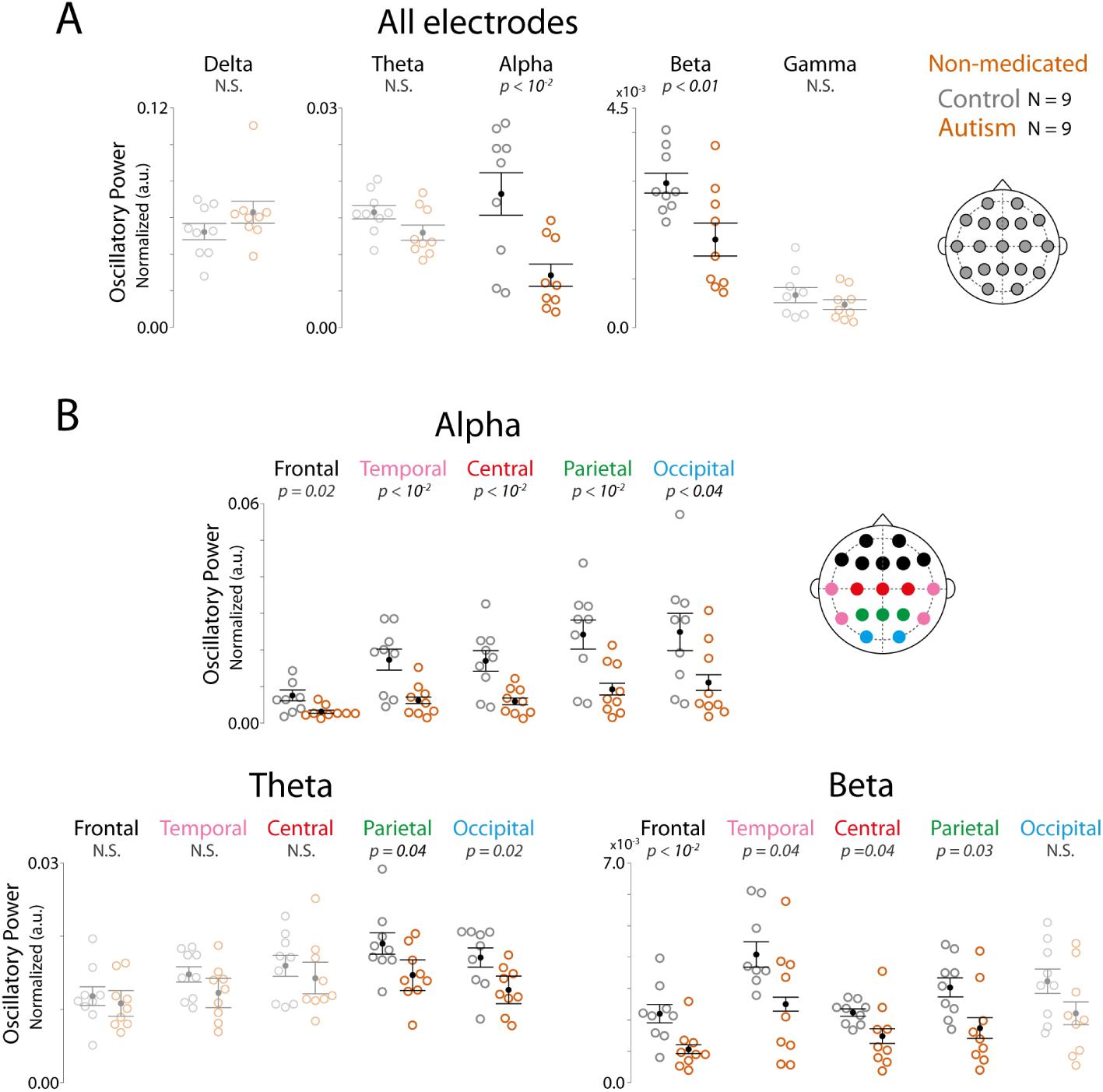
Full-band spectral power of ASD patients not taking any psychotropic medication reveals the alpha band as the most reliable biomarker for ASD. Same conventions as in Fig. 3. (A) Open circles represent the spectral power averaged within a certain frequency band for one subject across all electrodes. Filled dots represent the spectral power further averaged across all subjects. The associated thin line segments correspond to the standard errors of the mean (SEM). Control and ASD subjects are shown in gray and orange, respectively (N=9 each). Control vs. autism pairs exhibiting significant differences in their means (p<0.05) are highlighted, while those pairs that do not exhibit a significant difference are shown in pale colors. (B) Same conventions as in Panel A, but the results are broken down for the five brain regions studied (i.e., occipital, parietal, central, temporal and frontal regions). Results for the delta and gamma bands are not shown because they yielded no significant differences between control vs. ASD (see Figs 4 and 5). The frequency bands were defined as follows: delta (1–4 Hz), theta (4–8 Hz), alpha (8–12 Hz), beta (12–25 Hz) and gamma (25–85 Hz). The noise artifacts at and around 50 Hz and 60 Hz (i.e., the 45–75 Hz frequency segment) were excluded from the analysis.

**Table 3:** Non-medicated patients. Percentage change (% change) of EEG spectral power (autism relative to control) and its corresponding statistical significant (*p* value) for 5 frequency bands (delta, theta, alpha, beta and gamma) and 5 brain regions (frontal, temporal, central, parietal and occipital). Data from the subgroup of subjects not using psicotropic medication (ASD: N = 9, control: N = 9). The combined activity recorded from all electrodes (’All regions’) is also shown. Statistically significant changes are depicted in bold.

|  | Frequency Band |  |  |  |  |  |  |  |  |  |
|---|---|---|---|---|---|---|---|---|---|---|
|  | Delta |  | Theta |  | Alpha |  | Beta |  | Gamma |  |
| Brain region | % change | *p* | % change | *p* | % change | *p* | % change | *p* | % change | *p* |
| All regions | +20.32 | 0.20 | −19.72 | 0.07 | **−60.82** | **5.5 x 10<sup>-3</sup>** | **−39.05** | **1.4 x 10<sup>-2</sup>** | −29.13 | 0.34 |
| Frontal | +6.05 | 0.69 | −8.38 | 0.58 | **−58.86** | **1.5 x 10<sup>-2</sup>** | **−51.29** | **8.3 x 10<sup>-3</sup>** | −54.03 | 0.05 |
| Temporal | +37.75 | 0.07 | −17.13 | 0.15 | **−64.14** | **4.6 x 10<sup>-3</sup>** | **−38.73** | **4.7 x 10<sup>-2</sup>** | −24.33 | 0.52 |
| Central | +19.90 | 0.24 | −10.50 | 0.47 | **−65.01** | **3.4 x 10<sup>-3</sup>** | **−33.56** | **4.9 x 10<sup>-2</sup>** | −15.18 | 0.65 |
| Parietal | +22.87 | 0.21 | **−22.75** | **4.6 x 10<sup>-2</sup>** | **−64.14** | **4.6 x 10<sup>-3</sup>** | **−38.73** | **4.7 x 10<sup>-2</sup>** | −24.33 | 0.52 |
| Occipital | +21.84 | 0.30 | **−25.84** | **2.3 x 10<sup>-2</sup>** | **−55.50** | **4.6 x 10<sup>-2</sup>** | −31.54 | 0.12 | −25.99 | 0.53 |

Overall, our data shows that oscillatory dynamics in autism is marked by widespread power suppression encompassing frequencies ranging from ∼ 6 Hz to 20 Hz. Alpha, the frequency band in-between the 6 Hz to 20 Hz segment, is consistently suppressed in medicated and non-medicated subjects. ASD-associated theta suppression was mainly observed in posterior brain regions, while beta suppression was more strongly associated with frontal brain regions. Low theta (∼ 4 Hz to 6 Hz.) was the frequency band most affected by medication, even though future studies with a larger population of patients will be required to better access this result. Below, we discuss possible associations between weaker oscillations and the main behavioral and cognitive features associated with autism spectrum disorder.

## 4 Discussion

We recorded the resting state (eyes open) EEG signal of children and adolescents diagnosed with autism spectrum disorder (ASD). The results were compared with age- and gender-matched control subjects. Our results for the population including medicated and non-medicated subjects show a robust suppression of alpha and theta oscillations in all brain regions of ASD patients compared to controls. Beta oscillations were also suppressed, but the effect was more significantly observed in anterior brain regions. Notably, we were able to record from a subset of ASD patients with no history of psychotropic medication. We observed the same pattern of power suppression for all major brain regions, especially for the alpha frequency band (Fig. 1). Power in the delta and gamma frequency bands showed poor (if any) modulation due to ASD.

Here, we provide robust estimates of the raw power spectrum across frequency bands. This was possible due to three reasons. First, we used long EEG recording epochs, which provided spectra with high frequency resolution. Second, we used the multitaper method to compute all power spectra, which provided optimal spectral concentration, enabling us to better visualize the frequency peaks. Third, we excluded from our analysis all subjects that exhibited substantial muscle movement during the recordings. Together, these steps enabled reliable estimates of the raw power spectra computed from the brain activity of ASD patients.

Our data is unique in a few ways. First, we observe a selective ASD-associated power suppression for frequencies between 6 Hz and 20 Hz, encompassing the theta, alpha and beta bands. Notably, the neighboring bands on the lower (i.e., delta) and higher (i.e., gamma) parts of the spectrum underwent no significant power change. This localized spectral modulation reassures us that we are not observing a widespread oscillatory suppression in ASD, as was the case in Gabard-Durnam et al. [24] and Tierney et al. [35]. Second, the ASD-associated power modulation we observe is exclusively of suppressive nature. Other authors have described alpha and theta suppression in ASD but have concomitantly observed power enhancement for beta and/or gamma^27–32^. Third, we provide raw, continuous and high-resolution power spectra for the entire 1 to 85 Hz frequency range. This allows for a comprehensive analysis of power behavior, showing the robustness and consistency of the observed power suppression.

### 4.1 Functional significance of alpha suppression in autistic patients

Abnormal sensory processing affects 60% to 90% of autistic children^71–73^. For this reason, the DSM-5 includes sensory abnormalities as diagnostic criteria for autism. Auditory perception is reported to be the most impacted sensory modality (Dunn et al, 2002; Rogers et al, 2003; Wiggins et al, 2009), potentially leading to hyperacusis (reviewed in Danesh et al. [74]). Children and adults with ASD also show hyperreactivity to touch and abnormal pressure pain perception^75–79^. Other patients may show avoidance to specific odors^80^ or tastes^81,82^ or to excessive light^83^. We believe that the hyperreactivity that autistic subjects commonly show to incoming sensory stimuli^84–87^ may be the result of a lack of inhibitory control enabled by alpha. Indeed, alpha oscillations are thought to filter out irrelevant information, actively suppressing noise and distractors during perceptual processing^52^.

Alpha oscillations have also been implicated in feedback control mechanisms. Recent studies have shown that alpha waves originate in higher-order cortical regions and propagate in a feedback manner to early sensory areas^50,51^. These findings are compatible with the proposal by Klimesch, Sauseng, and Hanslmayr [88] who argued that alpha, in contrast to other oscillations, enable active top-down processes, such as attention, perception, functional inhibition, and working memory. This top-down alpha coordination may operate alongside the alpha inhibitory (i.e., “idling”) role mentioned earlier. One possibility is that the precise rhythmic timing of alpha oscillations enables selective patterns of inhibitory control and neuronal activation capable of actively coordinating downstream networks^52,88^.

A current hypothesis is that deficits in predictive coding can account for much of the perceptual and behavioral deficits associated with ASD^89–92^. This theory states that the brain is constantly generating a model of the world^93,94^. Some authors believe that the internal model shapes reality by restricting the influence of incoming sensory stimulation on conscious perception. In autism, this attenuation is compromised in a way that the external world becomes all “too real” and overwhelming^89^. We hypothesize that alpha oscillations are key to predictive coding in autism, since it embodies the feedback mechanism that conveys the expectations generated by the model to lower-level cortical regions. Therefore, the alpha suppression we observe in our data may be a key underlying mechanism leading to atypical perception in autism.

### 4.2 Functional significance of theta suppression in autistic patients

In our data, the prevalence of ASD-associated theta suppression was second only to the widespread alpha suppression described above. In our general population (medicated and non-medicated subjects), we observed significant theta power suppression for all brain regions. In our non-medicated subgroup, all regions showed a trend for suppression, but only the more posterior parietal and occipital regions showed a significant suppression.

Several studies report lower working memory performance in autism^95–99^. Interestingly, theta oscillations in the frontal and parietal cortices have been associated with working memory in neurotypical subjects^100–104^. Frontal theta has been hypothesized to orchestrate the maintenance of working memory content in posterior parietal regions^105–107^. In our data, we observe a more prominent ASD-associated deficit of theta oscillations in posterior (i.e., parietal and occipital) as compared to anterior (i.e., frontal) brain regions. Therefore, our data suggest a disruption of working memory representation, rather than a deficit in working memory orchestration across brain regions in autism. Regardless of which component is affected, working memory disruption in autism can account for some of its secondary deficits, such as learning, behavior regulation, cognitive flexibility, abstract thinking, sustained attention and focus^108^.

### 4.3 Functional significance of beta suppression in autistic patients

In our results, ASD-associated beta-suppression was more robustly observed in the frontal and temporal electrodes. Notably, we observe a significant beta suppression in the frontal region that extends up to 25 Hz, and which is also evident in the non-medicated subgroup. Other authors have also observed beta suppression in autism^109^. Beta oscillations in the frontal cortex have been associated with several aspects of executive function^110,111^, which is impaired in autism^112,113^. This includes deficits in planning, inhibitory control, impulse control, emotional self-control, mental flexibility, and autoregulation^114^. Inhibitory control may be one of the most critical components of frontal executive function underlying atypical behavior in ASD^115^. In our data, temporal electrodes exhibited stronger beta oscillations compared to all other brain regions. Özgören, Başar-Eroğlu, and Başar [116], recording the EEG of neurotypical subjects, observed a significant role for beta waves in face recognition. Even though their work did not focus on the contribution of the temporal electrodes, this region is classically known for its role in object and face processing^117^. Importantly, face processing^118–121^, the interpretation of facial emotions^122^ and holistic face recognition^123^ are impaired in autism. Therefore, beta desynchronization may disorganize the temporal and frontal networks involved in face processing and inhibitory control, respectively.

### 4.4 Impact of medication in the spectral profile of autistic patients

Medication use was carefully monitored in our study. Approximately 37% of our patients never used psychotropic drugs, while around 63% were on psychotropic medications at the time of the study. Of those patients on medication, 80% were on risperidone, an atypical antipsychotic medication that acts as an antagonist of both dopamine and serotonin neurotransmitters.

Work from several research groups have found that risperidone acts to increase the spectral power of a wide range of neuronal oscillations. Some found an enhancement of theta, alpha, and beta waves^124^, others of alpha and beta waves^68^, and others of alpha only^125,126^. Notably, these are the frequency bands that we found to be significantly suppressed in autism.

Above, we discussed possible associations between the oscillatory suppression we observed, and the corresponding behavioral and cognitive traits associated with autism. These associations remain speculative for now. However, they offer hypothesis-driven proposals for targeted behavioral paradigms in association with EEG recordings. They also offer a framework for testing the therapeutic benefits of transcranial electric or magnetic stimulation^127^ tailored to the frequency bands which are suppressed in ASD compared to neurotypical subjects. Notably, while attempting to characterize a possible EEG-based biomarker for autism, we may be able to simultaneously shed light into some of the underlying brain processes that are reshaped during non-neurotypical brain development.

## Statements and Declarations

The authors declare that this work was conducted in the absence of any commercial or financial relationships that could be construed as a potential conflict of interest. Additionally, we did not employ any AI-powered tools (like ChatGPT, LLMs, or similar software) for data analysis, writing, or content generation. This work was done entirely by human effort and the tools mentioned in the text.

## Acknowledgements

This study was supported by the Financiadora de Estudos e Projetos (FINEP) (Grant Number: 01.17.003.01), the Fundação Carlos Chagas Filho de Amparo à Pesquisa do Estado do Rio de Janeiro (FAPERJ) (Grant Numbers: E-26/211.258/2019, E-26/201.357/2021 and E-26/200.360/2026), the IDOR/Pioneer Science Initiative, and the Serrapilheira Institute (Grant Number: Serra-1709-17523). We would like to pay tribute to Amira C. M. Fiqueiras (*in memoriam*) for her invaluable clinical contribution during the early phases of this work.

## References

[1] American Psychiatric Association. Diagnostic and statistical manual of mental disorders. 5th ed., text rev. American Psychiatric Association, 2022. doi: 10.1176/appi.books.9780890425787. url: https://doi.org/10.1176/appi.books.9780890425787.

[2] F. Dellapiazza et al. “Links between sensory processing, adaptive behaviours, and attention in children with autism spectrum disorder: A systematic review”. In: Psychiatry Research 270 (2018), pp. 78–88. doi: 10.1016/j.psychres.2018.09.023. url: https://doi.org/10.1016/j.psychres.2018.09.023.

[3] F. Apicella, V. Costanzo, and G Purpura. “Are early visual behavior impairments involved in the onset of autism spectrum disorders? Insights for early diagnosis and intervention”. In: European Journal of Pediatrics 179.2 (2020), pp. 225–234. doi: 10.1007/s00431-019-03562-x. url: https://doi.org/10.1007/s00431-019-03562-x.

[4] E. Masini et al. “An overview of the main genetic, epigenetic and environmental factors involved in autism spectrum disorder focusing on synaptic activity”. In: International Journal of Molecular Sciences 21.21 (2020). Article 8290. doi: 10.3390/ijms21218290. url: https://doi.org/10.3390/ijms21218290.

[5] M. D. Bauman et al. “Activation of the maternal immune system during pregnancy alters behavioral development of rhesus monkey offspring”. In: Biological Psychiatry 75.4 (2014), pp. 332–341. doi: 10.1016/j.biopsych.2013.06.025. url: https://doi.org/10.1016/j.biopsych.2013.06.025.

[6] A. Meltzer and J. Van de Water. “The role of the immune system in autism spectrum disorder”. In: Neuropsychopharmacology 42.1 (2017), pp. 284–298. doi: 10.1038/npp.2016.158. url: https://doi.org/10.1038/npp.2016.158.

[7] Y. Feng et al. “Diffusion trajectory of atypical morphological development in autism spectrum disorder”. In: Communications Biology 8.1 (2025). Article 1476. doi: 10.1038/s42003-025-07612-4. url: https://doi.org/10.1038/s42003-025-07612-4.

[8] I. Voineagu et al. “Transcriptomic analysis of autistic brain reveals convergent molecular pathology”. In: Nature 474.7351 (2011), pp. 380–384. doi: 10.1038/nature10323. url: https://doi.org/10.1038/nature10323.

[9] L. E. Mash et al. “Atypical relationships between spontaneous EEG and fMRI activity in autism”. In: Brain Connectivity 10.1 (2020), pp. 18–28. doi: 10.1089/brain.2019.0705. url: https://doi.org/10.1089/brain.2019.0705.

[10] A. S. Chan and W. W Leung. “Differentiating autistic children with quantitative encephalography: A 3-month longitudinal study”. In: Journal of Child Neurology 21.5 (2006), pp. 391–399. doi: 10.1177/08830738060210050501. url: https://doi.org/10.1177/08830738060210050501.

[11] A. S. Chan, S. L. Sze, and M. C Cheung. “Quantitative electroencephalographic profiles for children with autistic spectrum disorder”. In: Neuropsychology 21.1 (2007), pp. 74–81. doi: 10.1037/0894-4105.21.1.74. url: https://doi.org/10.1037/0894-4105.21.1.74.

[12] R. Coben et al. “EEG power and coherence in autistic spectrum disorder”. In: Clinical Neurophysiology 119.5 (2008), pp. 1002–1009. doi: 10.1016/j.clinph.2008.01.013. url: https://doi.org/10.1016/j.clinph.2008.01.013.

[13] L. Cornew et al. “Resting-state oscillatory activity in autism spectrum disorders”. In: Journal of Autism and Developmental Disorders 42.9 (2012), pp. 1884–1894. doi: 10.1007/s10803-011-1431-6. url: https://doi.org/10.1007/s10803-011-1431-6.

[14] A. M. Daoust et al. “EEG spectral analysis of wakefulness and REM sleep in high functioning autistic spectrum disorders”. In: Clinical Neurophysiology 115.6 (2004), pp. 1368–1373. doi: 10.1016/j.clinph.2004.01.011. url: https://doi.org/10.1016/j.clinph.2004.01.011.

[15] H. Elhabashy et al. “Quantitative EEG in autistic children”. In: *The Egyptian Journal of Neurology*, Psychiatry and Neurosurgery 52.3 (2015), pp. 176–182. doi: 10.4103/1110-1083.162032. url: https://doi.org/10.4103/1110-1083.162032.

[16] N. J. Kozhushko et al. “Specificity of spontaneous EEG associated with different levels of cognitive and communicative dysfunctions in children”. In: International Journal of Psychophysiology 128 (2018), pp. 22–30. doi: 10.1016/j.ijpsycho.2018.03.016. url: https://doi.org/10.1016/j.ijpsycho.2018.03.016.

[17] N. Pop-Jordanova et al. “QEEG characteristics and spectrum weighted frequency for children diagnosed as autistic spectrum disorder”. In: Nonlinear Biomedical Physics 4.1 (2010). Article 4. doi: 10.1186/1753-4631-4-4. url: https://doi.org/10.1186/1753-4631-4-4.

[18] T. A. Stroganova et al. “Abnormal EEG lateralization in boys with autism”. In: Clinical Neurophysiology 118.8 (2007), pp. 1842–1854. doi: 10.1016/j.clinph.2007.05.004. url: https://doi.org/10.1016/j.clinph.2007.05.004.

[19] J. Wang et al. “Resting state EEG abnormalities in autism spectrum disorders”. In: Journal of Neurodevelopmental Disorders 5.1 (2013). Article 24. doi: 10.1186/1866-1955-5-24. url: https://doi.org/10.1186/1866-1955-5-24.

[20] B. Y. Angulo-Ruiz et al. “Linear and non-linear analyses of EEG in a group of ASD children during resting state condition”. In: Brain Topography 36.5 (2023), pp. 736–749. doi: 10.1007/s10548-023-00976-7. url: https://doi.org/10.1007/s10548-023-00976-7.

[21] M. Bink et al. “EEG theta and beta power spectra in adolescents with ADHD versus adolescents with ASD + ADHD”. In: European Child & Adolescent Psychiatry 24.8 (2015), pp. 873–886. doi: 10.1007/s00787-014-0632-x. url: https://doi.org/10.1007/s00787-014-0632-x.

[22] G. Dawson, S. Warrenburg, and P Fuller. “Cerebral lateralization in individuals diagnosed as autistic in early childhood”. In: Brain and Language 15.2 (1982), pp. 353–368. doi: 10.1016/0093-934X(82)90065-8. url: https://doi.org/10.1016/0093-934X(82)90065-8.

[23] G. Dawson et al. “Subgroups of autistic children based on social behavior display distinct patterns of brain activity”. In: Journal of Abnormal Child Psychology 23.5 (1995), pp. 569–583. doi: 10.1007/BF01447662. url: https://doi.org/10.1007/BF01447662.

[24] L. J. Gabard-Durnam et al. “Longitudinal EEG power in the first postnatal year differentiates autism outcomes”. In: Nature Communications 10.1 (2019). Article 4188. doi: 10.1038/s41467-019-12202-9. url: https://doi.org/10.1038/s41467-019-12202-9.

[25] S. Gulati et al. Comparison of blood heavy metal levels and quantitative EEG correlates in children with autism spectrum disorder aged 3–12 years and typically developing children: A cross sectional study. Conference presentation. 19th Annual Meeting of the International Society for Autism Research (INSAR), Virtual. June 2020.

[26] T. Hornung et al. “Dopaminergic hypoactivity and reduced theta-band power in autism spectrum disorder: A resting state EEG study”. In: International Journal of Psychophysiology 146 (2019), pp. 101–106. doi: 10.1016/j.ijpsycho.2019.09.006. url: https://doi.org/10.1016/j.ijpsycho.2019.09.006.

[27] J. Kang et al. “The identification of children with autism spectrum disorder by SVM approach on EEG and eye-tracking data”. In: Computers in Biology and Medicine 120 (2020). Article 103722. doi: 10.1016/j.compbiomed.2020.103722. url: https://doi.org/10.1016/j.compbiomed.2020.103722.

[28] E. A. Lushchekina et al. “Comparative studies of EEG theta and gamma rhythms in normal children and children with early childhood autism”. In: Neuroscience and Behavioral Physiology 44.8 (2014), pp. 902–908. doi: 10.1007/s11055-014-9999-5. url: https://doi.org/10.1007/s11055-014-9999-5.

[29] E. A. Lushchekina et al. “Interhemispheric differences in the spectral power and coherence of EEG rhythms in children with autism spectrum disorders”. In: Human Physiology 43.3 (2017), pp. 265–273. doi: 10.1134/S0362119717030113. url: https://doi.org/10.1134/S0362119717030113.

[30] E. A. Luschekina, V. S. Luschekin, and V. B. Strelets. “EEG Spectral Power in Children with Autistic Spectrum Disorders: Heterogeneity of the Group”. In: Human Physiology 45.3 (2019), pp. 242–248. doi: 10.1134/S036211971902004X. url: https://doi.org/10.1134/S036211971902004X.

[31] C. Machado et al. “QEEG spectral and coherence assessment of autistic children in three different experimental conditions”. In: Journal of Autism and Developmental Disorders 45.2 (2015), pp. 406–424. doi: 10.1007/s10803-013-1909-5. url: https://doi.org/10.1007/s10803-013-1909-5.

[32] M. Murias et al. “Resting state cortical connectivity reflected in EEG coherence in individuals with autism”. In: Biological Psychiatry 62.3 (2007), pp. 270–273. doi: 10.1016/j.biopsych.2006.11.012. url: https://doi.org/10.1016/j.biopsych.2006.11.012.

[33] E. Neuhaus et al. “Resting state EEG in youth with ASD: Age, sex, and relation to phenotype”. In: Journal of Neurodevelopmental Disorders 13.1 (2021). Article 33. doi: 10.1186/s11689-021-09383-7. url: https://doi.org/10.1186/s11689-021-09383-7.

[34] E. Shephard et al. “Resting state neurophysiological activity patterns in young people with ASD, ADHD, and ASD + ADHD”. In: Journal of Autism and Developmental Disorders 48.1 (2018), pp. 110–122. doi: 10.1007/s10803-017-3301-4. url: https://doi.org/10.1007/s10803-017-3301-4.

[35] A. L. Tierney et al. “Developmental trajectories of resting EEG power: An endophenotype of autism spectrum disorder”. In: PLoS ONE 7.6 (2012). Article e39127. doi: 10.1371/journal.pone.0039127. url: https://doi.org/10.1371/journal.pone.0039127.

[36] D. S. Cantor et al. “Computerized EEG analyses of autistic children”. In: Journal of Autism and Developmental Disorders 16.2 (1986), pp. 169–187. doi: 10.1007/BF01531728. url: https://doi.org/10.1007/BF01531728.

[37] K. J. Mathewson et al. “EEG alpha power, coherence, and behavioral symptomatology in autism spectrum disorder”. In: Clinical Neurophysiology 123.9 (2012), pp. 1798–1809. doi: 10.1016/j.clinph.2012.02.061. url: https://doi.org/10.1016/j.clinph.2012.02.061.

[38] E. V. Orekhova et al. “Excess of high frequency electroencephalogram oscillations in boys with autism”. In: Biological Psychiatry 62.9 (2007), pp. 1022–1029. doi: 10.1016/j.biopsych.2006.12.029. url: https://doi.org/10.1016/j.biopsych.2006.12.029.

[39] A. Sheikhani et al. “Abnormalities of quantitative electroencephalography in children with Asperger disorder in various conditions”. In: Research in Autism Spectrum Disorders 3.2 (2009), pp. 538–546. doi: 10.1016/j.rasd.2008.11.002. url: https://doi.org/10.1016/j.rasd.2008.11.002.

[40] E. van Diessen et al. “Increased power of resting-state gamma oscillations in autism spectrum disorder detected by routine electroencephalography”. In: European Archives of Psychiatry and Clinical Neuroscience 265.6 (2015), pp. 537–540. doi: 10.1007/s00406-014-0527-3. url: https://doi.org/10.1007/s00406-014-0527-3.

[41] A. Bellato et al. “Atypical electrophysiological indices of eyes-open and eyes-closed resting-state in children and adolescents with ADHD and autism”. In: Brain Sciences 10.5 (2020). Article 272. doi: 10.3390/brainsci10050272. url: https://doi.org/10.3390/brainsci10050272.

[42] C. Distefano et al. “EEG data collection in children with ASD: The role of state in data quality and spectral power”. In: Research in Autism Spectrum Disorders 57 (2019), pp. 132–144. doi: 10.1016/j.rasd.2018.10.010. url: https://doi.org/10.1016/j.rasd.2018.10.010.

[43] J. Frohlich et al. “A quantitative electrophysiological biomarker of duplication 15q11.2-q13.1 syndrome”. In: PLoS ONE 11.12 (2016). Article e0167179. doi: 10.1371/journal.pone.0167179. url: https://doi.org/10.1371/journal.pone.0167179.

[44] P. J. Uhlhaas and W Singer. “What do disturbances in neural synchrony tell us about autism?” In: Biological Psychiatry 62.3 (2007), pp. 190–191. doi: 10.1016/j.biopsych.2007.05.023. url: https://doi.org/10.1016/j.biopsych.2007.05.023.

[45] P. J. Uhlhaas and W Singer. “Neuronal dynamics and neuropsychiatric disorders: Toward a translational paradigm for dysfunctional large-scale networks”. In: Neuron 75.6 (2012), pp. 963–980. doi: 10.1016/j.neuron.2012.09.004. url: https://doi.org/10.1016/j.neuron.2012.09.004.

[46] E. A. Lushchekina et al. “Characteristics of the spectral power of EEG rhythms in children with early childhood autism and their association with the development of different symptoms of schizophrenia”. In: Neuroscience and Behavioral Physiology 43.1 (2013), pp. 40–45. doi: 10.1007/s11055-012-9694-z. url: https://doi.org/10.1007/s11055-012-9694-z.

[47] E. A. Luschekina and V. B. Strelets. “Comparative analysis of autism spectrum disorders and schizophrenia in childhood”. In: Human Physiology 46.1 (2020), pp. 94–103. doi: 10.1134/S0362119720010119. url: https://doi.org/10.1134/S0362119720010119.

[48] C. R. Maxwell et al. “Atypical laterality of resting gamma oscillations in autism spectrum disorders”. In: Journal of Autism and Developmental Disorders 45.2 (2013), pp. 292–297. doi: 10.1007/s10803-013-1842-7. url: https://doi.org/10.1007/s10803-013-1842-7.

[49] M Steriade. “Corticothalamic resonance, states of vigilance and mentation”. In: Neuroscience 101.2 (2000), pp. 243–276. doi: 10.1016/S0306-4522(00)00353-5. url: https://doi.org/10.1016/S0306-4522(00)00353-5.

[50] M. Halgren et al. “The generation and propagation of the human alpha rhythm”. In: Proceedings of the National Academy of Sciences 116.47 (2019), pp. 23772–23782. doi: 10.1073/pnas.1913092116. url: https://doi.org/10.1073/pnas.1913092116.

[51] T. van Kerkoerle et al. “Alpha and gamma oscillations characterize feedback and feed-forward processing in monkey visual cortex”. In: Proceedings of the National Academy of Sciences 111.40 (2014), pp. 14332–14341. doi: 10.1073/pnas.1402773111. url: https://doi.org/10.1073/pnas.1402773111.

[52] W Klimesch. “Alpha-band oscillations, attention, and controlled access to stored information”. In: Trends in Cognitive Sciences 16.12 (2012), pp. 606–617. doi: 10.1016/j.tics.2012.10.007. url: https://doi.org/10.1016/j.tics.2012.10.007.

[53] W. Klimesch et al. “Induced alpha band power changes in the human EEG and attention”. In: Neuroscience Letters 244.2 (1998), pp. 73–76. doi: 10.1016/S0304-3940(98)00122-0. url: https://doi.org/10.1016/S0304-3940(98)00122-0.

[54] J. Mo, C. E. Schroeder, and M Ding. “Attentional modulation of alpha oscillations in macaque inferotemporal cortex”. In: The Journal of Neuroscience 31.3 (2011), pp. 878–882. doi: 10.1523/JNEUROSCI.5295-10.2011. url: https://doi.org/10.1523/JNEUROSCI.5295-10.2011.

[55] S. Palva and J. M Palva. “Functional roles of alpha-band phase synchronization in local and large-scale cortical networks”. In: Frontiers in Psychology 2 (2011). Article 204. doi: 10.3389/fpsyg.2011.00204. url: https://doi.org/10.3389/fpsyg.2011.00204.

[56] K. M. Burner et al. Frontal asymmetry and temperament in young high functioning children with autism spectrum disorders. Poster presentation. 10th International Meeting for Autism Research (IMFAR), San Diego, CA, United States. May 2011.

[57] J. C. Edgar et al. “Resting-state alpha in autism spectrum disorder and alpha associations with thalamic volume”. In: Journal of Autism and Developmental Disorders 45.3 (2015), pp. 795–804. doi: 10.1007/s10803-014-2236-1. url: https://doi.org/10.1007/s10803-014-2236-1.

[58] S. K. Sutton et al. “Resting cortical brain activity and social behavior in higher functioning children with autism”. In: Journal of Child Psychology and Psychiatry 46.2 (2005), pp. 211–222. doi: 10.1111/j.1469-7610.2004.00354.x. url: https://doi.org/10.1111/j.1469-7610.2004.00354.x.

[59] H. Bruining et al. “Measurement of excitation-inhibition ratio in autism spectrum disorder using critical brain dynamics”. In: Scientific Reports 10.1 (2020). Article 9195. doi: 10.1038/s41598-020-65500-4. url: https://doi.org/10.1038/s41598-020-65500-4.

[60] A. Dickinson et al. “Peak alpha frequency is a neural marker of cognitive function across the autism spectrum”. In: European Journal of Neuroscience 47.6 (2018), pp. 643–651. doi: 10.1111/ejn.13845. url: https://doi.org/10.1111/ejn.13845.

[61] L. J. Gabard-Durnam et al. “Alpha asymmetry in infants at risk for autism spectrum disorders”. In: Journal of Autism and Developmental Disorders 45.2 (2015), pp. 473–480. doi: 10.1007/s10803-013-1926-4. url: https://doi.org/10.1007/s10803-013-1926-4.

[62] B. Keehn et al. “Autism, attention, and alpha oscillations: An electrophysiological study of attentional capture”. In: Biological Psychiatry: Cognitive Neuroscience and Neuroimaging 2.6 (2017), pp. 528–536. doi: 10.1016/j.bpsc.2017.06.006. url: https://doi.org/10.1016/j.bpsc.2017.06.006.

[63] S. Pierce et al. “Associations between sensory processing and electrophysiological and neurochemical measures in children with ASD: An EEG-MRS study”. In: Journal of Neurodevelopmental Disorders 13.1 (2021). Article 5. doi: 10.1186/s11689-020-09353-3. url: https://doi.org/10.1186/s11689-020-09353-3.

[64] P. Wantzen et al. “EEG resting-state functional connectivity: Evidence for an imbalance of external/internal information integration in autism”. In: Journal of Neurodevelopmental Disorders 14.1 (2022). Article 47. doi: 10.1186/s11689-022-09458-7. url: https://doi.org/10.1186/s11689-022-09458-7.

[65] A. J. Dede et al. “Exploring EEG resting state differences in autism: Sparse findings from a large cohort”. In: Molecular Autism 16.1 (2025), pp. 1–13. doi: 10.1186/s13229-024-00632-y. url: https://doi.org/10.1186/s13229-024-00632-y.

[66] W. S. Neo et al. “Resting-state EEG power differences in autism spectrum disorder: A systematic review and meta-analysis”. In: Translational Psychiatry 13.1 (2023). Article 389. doi: 10.1038/s41398-023-02685-6. url: https://doi.org/10.1038/s41398-023-02685-6.

[67] S. Schwartz et al. “Electroencephalogram coherence patterns in autism: An updated review”. In: Pediatric Neurology 67 (2017), pp. 7–22. doi: 10.1016/j.pediatrneurol.2016.09.014. url: https://doi.org/10.1016/j.pediatrneurol.2016.09.014.

[68] T. Ozaki et al. “Quantitative resting state electroencephalography in patients with schizophrenia spectrum disorders treated with strict monotherapy using atypical antipsychotics”. In: Clinical Psychopharmacology and Neuroscience 19.2 (2021), pp. 313–322. doi: 10.9758/cpn.2021.19.2.313. url: https://doi.org/10.9758/cpn.2021.19.2.313.

[69] P. Mitra and H Bokil. Observed brain dynamics. Oxford University Press, 2008. doi: 10.1093/acprof:oso/9780195178081.001.0001. url: 10.1093/acprof:oso/9780195178081.001.0001.

[70] D. J Thomson. “Spectrum estimation and harmonic analysis”. In: Proceedings of the IEEE 70.9 (1982), pp. 1055–1096. doi: 10.1109/PROC.1982.12433. url: https://doi.org/10.1109/PROC.1982.12433.

[71] A. Ben-Sasson et al. “Update of a meta-analysis of sensory symptoms in ASD: A new decade of research”. In: Journal of Autism and Developmental Disorders 49.12 (2019), pp. 4974–4996. doi: 10.1007/s10803-019-04180-0. url: https://doi.org/10.1007/s10803-019-04180-0.

[72] K. R. Parmar et al. “Visual sensory experiences from the viewpoint of autistic adults”. In: Frontiers in Psychology 12 (2021). Article 633037. doi: 10.3389/fpsyg.2021.633037. url: https://doi.org/10.3389/fpsyg.2021.633037.

[73] T. Tavassoli et al. “Measuring sensory reactivity in autism spectrum disorder: Application and simplification of a clinician-administered sensory observation scale”. In: Journal of Autism and Developmental Disorders 46.1 (2016), pp. 287–293. doi: 10.1007/s10803-015-2578-3. url: https://doi.org/10.1007/s10803-015-2578-3.

[74] A. A. Danesh et al. “Hyperacusis in autism spectrum disorders”. In: Audiology Research 11.4 (2021), pp. 547–556. doi: 10.3390/audiolres11040050. url: https://doi.org/10.3390/audiolres11040050.

[75] S. R. Leekam et al. “Describing the sensory abnormalities of children and adults with autism”. In: Journal of Autism and Developmental Disorders 37.5 (2007), pp. 894–910.doi: 10.1007/s10803-006-0218-7. url: https://doi.org/10.1007/s10803-006-0218-7.

[76] M. D. Kaiser et al. “Brain mechanisms for processing affective (and nonaffective) touch are atypical in autism”. In: Cerebral Cortex 26.6 (2016), pp. 2705–2714. doi: 10.1093/cercor/bhv125. url: https://doi.org/10.1093/cercor/bhv125.

[77] I. Riquelme, S. M. Hatem, and P Montoya. “Abnormal pressure pain, touch sensitivity, proprioception, and manual dexterity in children with autism spectrum disorders”. In: Neural Plasticity 2016 (2016). Article 1723401. doi: 10.1155/2016/1723401. url: https://doi.org/10.1155/2016/1723401.

[78] T. Tavassoli et al. “Altered tactile processing in children with autism spectrum disorder”. In: Autism Research 9.6 (2016), pp. 616–620. doi: 10.1002/aur.1563. url: https://doi.org/10.1002/aur.1563.

[79] S. Vaughan et al. “A quantitative sensory testing approach to pain in autism spectrum disorders”. In: Journal of Autism and Developmental Disorders 50.5 (2020), pp. 1607–1620. doi: 10.1007/s10803-019-04331-3. url: https://doi.org/10.1007/s10803-019-04331-3.

[80] A. Tonacci et al. “Olfaction in autism spectrum disorders: A systematic review”. In: Child Neuropsychology 23.1 (2017), pp. 1–25. doi: 10.1080/09297049.2015.1081678. url: https://doi.org/10.1080/09297049.2015.1081678.

[81] S. J. Rogers, S. Hepburn, and E Wehner. “Parent reports of sensory symptoms in toddlers with autism and those with other developmental disorders”. In: Journal of Autism and Developmental Disorders 33.6 (2003), pp. 631–642. doi: 10.1023/B:JADD.0000006000.38991.a7. url:https://doi.org/10.1023/B:JADD.0000006000.38991.a7.

[82] L. D. Wiggins et al. “Brief report: Sensory abnormalities as distinguishing symptoms of autism spectrum disorders in young children”. In: Journal of Autism and Developmental Disorders 39.7 (2009), pp. 1087–1091. doi: 10.1007/s10803-009-0711-x. url: https://doi.org/10.1007/s10803-009-0711-x.

[83] T. Tavassoli et al. “Sensory over-responsivity in adults with autism spectrum conditions”. In: Autism 18.4 (2014), pp. 428–432. doi: 10.1177/1362361313477246. url: https://doi.org/10.1177/1362361313477246.

[84] S. Baron-Cohen et al. “Talent in autism: Hyper-systemizing, hyper-attention to detail and sensory hypersensitivity”. In: Philosophical Transactions of the Royal Society B: Biological Sciences 364.1522 (2009), pp. 1377–1383. doi: 10.1098/rstb.2008.0337. url: https://doi.org/10.1098/rstb.2008.0337.

[85] S. E. Christ et al. “Evidence for selective inhibitory impairment in individuals with autism spectrum disorder”. In: Neuropsychology 25.6 (2011), pp. 690–701. doi: 10.1037/a0024227. url: https://doi.org/10.1037/a0024227.

[86] A. M. Remington, J. G. Swettenham, and N Lavie. “Lightening the load: Perceptual load impairs visual detection in typical adults but not in autism”. In: Journal of Abnormal Psychology 121.2 (2012), pp. 544–551. doi: 10.1037/a0025341. url: https://doi.org/10.1037/a0025341.

[87] C. E. Robertson and S Baron-Cohen. “Sensory perception in autism”. In: Nature Reviews Neuroscience 18.11 (2017), pp. 671–684. doi: 10.1038/nrn.2017.112. url: https://doi.org/10.1038/nrn.2017.112.

[88] W. Klimesch, P. Sauseng, and S Hanslmayr. “EEG alpha oscillations: The inhibition-timing hypothesis”. In: Brain Research Reviews 53.1 (2007), pp. 63–88. doi: 10 . 1016/j.brainresrev.2006.06.003. url: https://doi.org/10.1016/j.brainresrev.2006.06.003.

[89] E. Pellicano and D Burr. “When the world becomes ’too real’: A Bayesian explanation of autistic perception”. In: Trends in Cognitive Sciences 16.10 (2012), pp. 504–510. doi: 10.1016/j.tics.2012.08.009. url: https://doi.org/10.1016/j.tics.2012.08.009.

[90] R. P. Lawson, G. Rees, and K. J Friston. “An aberrant precision account of autism”. In: Frontiers in Human Neuroscience 8 (2014). Article 302. doi: 10.3389/fnhum.2014.00302. url: https://doi.org/10.3389/fnhum.2014.00302.

[91] P. Sinha et al. “Autism as a disorder of prediction”. In: Proceedings of the National Academy of Sciences 111.42 (2014), pp. 15220–15225. doi: 10.1073/pnas.1416797111. url: https://doi.org/10.1073/pnas.1416797111.

[92] S. Van de Cruys et al. “Precise minds in uncertain worlds: Predictive coding in autism”. In: Psychological Review 121.4 (2014), pp. 649–675. doi: 10.1037/a0037665. url: https://doi.org/10.1037/a0037665.

[93] R. P. Rao and D. H Ballard. “Predictive coding in the visual cortex: A functional interpretation of some extra-classical receptive-field effects”. In: Nature Neuroscience 2.1 (1999), pp. 79–87. doi: 10.1038/4580. url: https://doi.org/10.1038/4580.

[94] D. M. Wolpert, Z. Ghahramani, and M. I Jordan. “An internal model for sensorimotor integration”. In: Science 269.5232 (1995), pp. 1880–1882. doi: 10.1126/science.7569931. url: https://doi.org/10.1126/science.7569931.

[95] P. Desaunay et al. “Memory in autism spectrum disorder: A meta-analysis of experimental studies”. In: Psychological Bulletin 146.5 (2020), pp. 377–410. doi: 10.1037/bul0000225. url: https://doi.org/10.1037/bul0000225.

[96] A. Habib et al. “A meta-analysis of working memory in individuals with autism spectrum disorders”. In: PLoS ONE 14.4 (2019). Article e0216198. doi: 10.1371/journal.pone.0216198. url: https://doi.org/10.1371/journal.pone.0216198.

[97] J. Larrain-Valenzuela et al. “Theta and alpha oscillation impairments in autistic spectrum disorder reflect working memory deficit”. In: Scientific Reports 7.1 (2017). Article 14328. doi: 10.1038/s41598-017-14611-8. url: https://doi.org/10.1038/s41598-017-14611-8.

[98] J. Liu et al. “Replicable patterns of memory impairments in children with autism and their links to hyperconnected brain circuits”. In: Biological Psychiatry: Cognitive Neuroscience and Neuroimaging 8.11 (2023), pp. 1113–1123. doi: 10.1016/j.bpsc.2023.05.006. url: https://doi.org/10.1016/j.bpsc.2023.05.006.

[99] M. Zhang et al. “Exploring the spatial working memory and visual perception in children with autism spectrum disorder and general population with high autism-like traits”. In: PLoS ONE 15.7 (2020). Article e0235552. doi: 10.1371/journal.pone.0235552. url: https://doi.org/10.1371/journal.pone.0235552.

[100] N. A. Herweg, E. A. Solomon, and M. J Kahana. “Theta oscillations in human memory”. In: Trends in Cognitive Sciences 24.3 (2020), pp. 208–227. doi: 10.1016/j.tics.2019.12.006. url: https://doi.org/10.1016/j.tics.2019.12.006.

[101] O. Jensen and C. D Tesche. “Frontal theta activity in humans increases with memory load in a working memory task”. In: European Journal of Neuroscience 15.8 (2002), pp. 1395–1399. doi: 10.1046/j.1460-9568.2002.01975.x. url: https://doi.org/10.1046/j.1460-9568.2002.01975.x.

[102] S. Raghavachari et al. “Gating of human theta oscillations by a working memory task”. In: The Journal of Neuroscience 21.9 (2001), pp. 3175–3183. doi: 10.1523/JNEUROSCI.21-09-03175.2001. url: https://doi.org/10.1523/JNEUROSCI.21-09-03175.2001.

[103] J. Riddle et al. “Causal evidence for a role of theta and alpha oscillations in the control of working memory”. In: Current Biology 30.9 (2020), pp. 1748–1754. doi: 10.1016/j.cub.2020.02.065. url: https://doi.org/10.1016/j.cub.2020.02.065.

[104] F. Roux and P. J Uhlhaas. “Working memory and neural oscillations: Alpha–gamma versus theta–gamma codes for distinct WM information?” In: Trends in Cognitive Sciences 18.1 (2014), pp. 16–25. doi: 10.1016/j.tics.2013.10.010. url: https://doi.org/10.1016/j.tics.2013.10.010.

[105] T. Imaruoka, J. Saiki, and S Miyauchi. “Maintaining coherence of dynamic objects requires coordination of neural systems extended from anterior frontal to posterior parietal brain cortices”. In: NeuroImage 26.1 (2005), pp. 277–284. doi: 10.1016/j.neuroimage.2005.01.042. url: https://doi.org/10.1016/j.neuroimage.2005.01.042.

[106] O. Ratcliffe, K. Shapiro, and B. P Staresina. “Fronto-medial theta coordinates posterior maintenance of working memory content”. In: Current Biology 32.10 (2022), pp. 2121–2129. doi: 10.1016/j.cub.2022.03.045. url: https://doi.org/10.1016/j.cub.2022.03.045.

[107] V. Yuk et al. “Frontoparietal network connectivity during an N-back task in adults with autism spectrum disorder”. In: Frontiers in Psychiatry 11 (2020). Article 551808. doi: 10.3389/fpsyt.2020.551808. url: https://doi.org/10.3389/fpsyt.2020.551808.

[108] S. Kercood et al. “Working memory and autism: A review of literature”. In: Research in Autism Spectrum Disorders 8.10 (2014), pp. 1316–1332. doi: 10.1016/j.rasd.2014.06.011. url: https://doi.org/10.1016/j.rasd.2014.06.011.

[109] J. Castelhano et al. “Stimulus dependent neural oscillatory patterns show reliable statistical identification of autism spectrum disorder in a face perceptual decision task”. In: Clinical Neurophysiology 129.5 (2018), pp. 981–989. doi: 10.1016/j.clinph.2018.01.072. url: https://doi.org/10.1016/j.clinph.2018.01.072.

[110] Y. M. Fonken et al. “Frontal and motor cortex contributions to response inhibition: Evidence from electrocorticography”. In: Journal of Neurophysiology 115.4 (2016), pp. 2224–2236. doi: 10.1152/jn.00708.2015. url: https://doi.org/10.1152/jn.00708.2015.

[111] J. Wagner et al. “Establishing a right frontal beta signature for stopping action in scalp EEG: Implications for testing inhibitory control in other task contexts”. In: Journal of Cognitive Neuroscience 30.1 (2018), pp. 107–118. doi: 10.1162/jocn_a_01183. url: https://doi.org/10.1162/jocn_a_01183.

[112] B. A. Corbett et al. “Examining executive functioning in children with autism spectrum disorder, attention deficit hyperactivity disorder and typical development”. In: Psychiatry Research 166.2–3 (2009), pp. 210–222. doi: 10.1016/j.psychres.2008.02.005. url: https://doi.org/10.1016/j.psychres.2008.02.005.

[113] E. A. Demetriou et al. “Autism spectrum disorders: A meta-analysis of executive function”. In: Molecular Psychiatry 23.5 (2018), pp. 1198–1204. doi: 10.1038/mp.2017.75. url: https://doi.org/10.1038/mp.2017.75.

[114] E. A. Demetriou, M. M. DeMayo, and A. J Guastella. “Executive function in autism spectrum disorder: History, theoretical models, empirical findings, and potential as an endophenotype”. In: Frontiers in Psychiatry 10 (2019). Article 753. doi: 10.3389/fpsyt.2019.00753. url: https://doi.org/10.3389/fpsyt.2019.00753.

[115] S. E. Christ et al. “Inhibitory control in children with autism spectrum disorder”. In: Journal of Autism and Developmental Disorders 37.6 (2007), pp. 1155–1165. doi: 10.1007/s10803-006-0259-y. url: https://doi.org/10.1007/s10803-006-0259-y.

[116] M. Özgören, C. Başar-Eroğlu, and E. Başar. “Beta oscillations in face recognition”. In: International Journal of Psychophysiology 55.1 (2005), pp. 51–59. doi: 10.1016/j.ijpsycho.2004.06.005. url: https://doi.org/10.1016/j.ijpsycho.2004.06.005.

[117] C. G. Gross, C. E. Rocha-Miranda, and D. B Bender. “Visual properties of neurons in inferotemporal cortex of the macaque”. In: Journal of Neurophysiology 35.1 (1972), pp. 96–111. doi: 10.1152/jn.1972.35.1.96. url: https://doi.org/10.1152/jn.1972.35.1.96.

[118] J. Boucher, V. Lewis, and G Collis. “Familiar face and voice matching and recognition in children with autism”. In: Journal of Child Psychology and Psychiatry 39.2 (1998), pp. 171–181. doi: 10.1111/1469-7610.00311. url: https://doi.org/10.1111/1469-7610.00311.

[119] L. Ewing, E. Pellicano, and G Rhodes. “Reevaluating the selectivity of face-processing difficulties in children and adolescents with autism”. In: Journal of Experimental Child Psychology 115.2 (2013), pp. 342–355. doi: 10.1016/j.jecp.2012.12.009. url: https://doi.org/10.1016/j.jecp.2012.12.009.

[120] A. Klin et al. “A normed study of face recognition in autism and related disorders”. In: Journal of Autism and Developmental Disorders 29.6 (1999), pp. 499–508. doi: 10.1023/A:1022299818817. url: https://doi.org/10.1023/A:1022299818817.

[121] S. Wallace, M. Coleman, and A Bailey. “Face and object processing in autism spectrum disorders”. In: Autism Research 1.1 (2008), pp. 43–51. doi: 10.1002/aur.4. url: https://doi.org/10.1002/aur.4.

[122] K. A. Pelphrey et al. “Visual scanning of faces in autism”. In: Journal of Autism and Developmental Disorders 32.4 (2002), pp. 249–261. doi: 10.1023/A:1016374617369. url: https://doi.org/10.1023/A:1016374617369.

[123] P. P. Tavares et al. “Preserved face inversion effects in adults with autism spectrum disorder: An event-related potential study”. In: NeuroReport 27.8 (2016), pp. 587–592. doi: 10.1097/WNR.0000000000000588. url: https://doi.org/10.1097/WNR.0000000000000588.

[124] A. M. Hughes et al. “Electroencephalographic and psychomotor effects of chlorpromazine and risperidone relative to placebo in normal healthy volunteers”. In: British Journal of Clinical Pharmacology 48.3 (1999), pp. 323–330. doi: 10.1046/j.1365-2125.1999.00010.x. url: https://doi.org/10.1046/j.1365-2125.1999.00010.x.

[125] J. Hyun, M. J. Baik, and U. G Kang. “Effects of psychotropic drugs on quantitative EEG among patients with schizophrenia-spectrum disorders”. In: Clinical Psychopharmacology and Neuroscience 9.2 (2011), pp. 78–85. doi: 10.9758/cpn.2011.9.2.78. url: https://doi.org/10.9758/cpn.2011.9.2.78.

[126] M. McVoy and R Findling. “Child and adolescent psychopharmacology update”. In: Psychiatric Clinics of North America 32.1 (2009), pp. 111–133. doi: 10.1016/j.psc.2008.11.002. url: https://doi.org/10.1016/j.psc.2008.11.002.

[127] U. Ezedinma, P. Swierkowski, and S Fjaagesund. “Outcomes from individual alpha frequency guided repetitive transcranial magnetic stimulation in children with autism spectrum disorder–a retrospective chart review”. In: Child Psychiatry & Human Development 55.4 (2024), pp. 1010–1019. doi: 10.1007/s10578-022-01460-2. url: https://doi.org/10.1007/s10578-022-01460-2.

